# Identification and characterization of pleiotropic high-persistence mutations in the beta subunit of the bacterial RNA polymerase

**DOI:** 10.1101/2020.08.19.258475

**Authors:** Lev Ostrer, Yinduo Ji, Arkady Khodursky

**Affiliations:** Dept. of Biochemistry. Molecular Biology and Biophysics, and Biotechnology Institute; Department of Veterinary Biomedical Sciences, College of Veterinary Medicine, University of Minnesota, St. Paul, Minnesota

**Keywords:** Persistence, RNA polymerase, RpoB, stringent response, *E. coli*, MRSA, MTB, antibiotic resistance, ppGpp, rifampicin, fluoroquinolones, ciprofloxacin

## Abstract

Individual bacteria can escape killing by bactericidal antibiotics by becoming dormant. Such cells, also known as persisters, naturally occur in bacterial populations at a low frequency. Here we present the finding that antibiotic-resistance mutations in the *rpoB* gene, encoding the beta subunit of RNA polymerase, increase the frequency of persisters by orders of magnitude. Furthermore, we show that: i) the persistent state depends on the (p)ppGpp transcriptional program and not on (p)ppGpp itself; ii) the high persistence (*hip*) is associated with increased populational heterogeneity in transcription; iii) indole overproduction, caused by transcriptional changes in the *hip* mutants, explains 50-80% of the *hip* phenotype. We report that the analogous *rpoB* mutations occur frequently in clinical isolates of *Acinetobacter baumannii, Mycobacterium tuberculosis and Staphylococcus aureus*, and we demonstrate that one of those *rpoB* mutations causes high persistence in MRSA. We also show that the RpoB-associated *hip* phenotype can be reversed by inhibiting protein synthesis.

**Importance:** Persistence is an inevitable consequence of antibiotic usage. Although persistence is not a genetically heritable trait, here we demonstrate for the first time that antibiotic resistance, which is heritable, can promote persistence formation. Our finding that resistance to one antibiotic, rifampicin, can boost persistence to other antibiotics, such as ciprofloxacin and ampicillin, may help explain why certain chronic infections are particularly recalcitrant to antibiotic therapies. Out results also emphasize the need to assess the effects of combination antibiotic therapies on persistence.

## Introduction

Persisters make up a small subpopulation of metabolically slowed cells that are recalcitrant to many types of stress, including antibiotics. Persisters can be induced via changes in the environment or arise spontaneously. Environmental challenges such as starvation (1) and exposure to chemicals/antibiotics (2) can increase the persister frequency manifold; such an increase can be mediated by peptides and metabolites (3,4). For example, indole, a by-product of tryptophan metabolism, can contribute to a rapid accumulation of persister cells in early stationary cultures (5,6), likely by reversibly inhibiting cell growth and division (7). Persisters can also arise spontaneously due to a heterogeneity of physiological states among individual cells in bacterial populations (8,9). The extent of the heterogeneity can be increased by mutations that unbalance toxin-antitoxin activity in the cell, the insight stemming from the work of Moyed *et al*. showing that overexpression of HipA toxin in certain *hipA* mutants increases persistence by 10^3^ to 10^4^ -fold (10,11). With several dozen identified toxin-antitoxin systems, others such as *relE/relB* and *mazE/mazF* have been also shown to contribute to the formation of the persister cells (12,13). The toxin-induced dormant state, in which bacteria are recalcitrant to bactericidal antibiotics, is achieved by a reversible growth inhibition (13,14).

Whether induced via macroscopic environmental cues or spontaneously generated, the persistence phenotype most of the time is triggered through stress-response regulatory circuit. Several stress-response pathways were demonstrated to be involved in the induction of the persister state, including SOS, oxidative stress, and stringent response (2,5,15,16). Stringent response is normally triggered by starvation (17). In the absence of a key nutrient(s), enzymes RelA and SpoT are recruited to synthesize the alarmone (p)ppGpp (18,19), which in turn interacts with RNA polymerase to globally adjust transcription within the cell (20). Bacteria utilizes stringent response to reprogram their metabolism when transitioning from rich to poor nutritional sources (21,22). Many toxin-antitoxins act by “hijacking” the stringent response pathway, forcing cells to experience starvation in the presence of nutrients and consequently triggering the persister state (23,24). In addition to starvation and toxin-antitoxin action, stringent response can be constitutively activated by mutations in the RNA polymerase (25,26). Although stringent RNA polymerase mutations have been extensively studied (27), their role in persister formation has been overlooked, presumably because of their relatively slow-growth phenotype (25). Here we disentangle the growth phenotype from the persister phenotype and demonstrate that *rpoB* stringent mutations represent a common evolutionary path toward high persistence in different bacterial species. We also show that mutations in the RpoB switch region (28,29), which could be readily selected for at low concentrations of the quinolone ciprofloxacin, mimic ppGpp transcriptional response and, as a result, confer high persistence phenotype.

## Results

### High survival frequency is caused by rpoB mutations

We have selected *E. coli* MG1655 mutants with improved survival frequency by incrementally increasing concentration of ciprofloxacin. Among mutants that were resistant to 0.25 mg/L of ciprofloxacin, but not 0.5 mg/L, we identified two independent clones with dramatically elevated survival at 4 mg/L. Whole-genome sequencing revealed that each of the mutants contained only two mutations: a *gyrB* mutation (S464Y) and an *rpoB* mutation, one of the mutants carried a substitution in 1279 residue of RpoB (E–>A) and another in 1304 (M–>R). To determine the optimal concentration of ciprofloxacin for testing strains’ persistence, at 10x MIC or higher, we measured MIC and MBC of the parent and the mutants. The GyrB S464Y substitution caused an increase in MIC but not in survival frequency (Table 1, Fig. 1A). By contrast, the RpoB E1279A and RpoB M1304R substitutions resulted in an 1.5-2-fold increase in MIC and MBC (Table 1) while also increasing the survival frequency at 10x MIC by more than 100 times (Fig. 1A).

**Table 1.** MIC/MBC

| **Parental genotype** | **mg/L ciprofloxacin** |  |
| --- | --- | --- |
|  | **MIC** | **MBC** |
| *MG1655* | 0.015 | 0.033 |
| *MG1655 RpoB 1304* | 0.025 | 0.055 |
| *MG1655 RpoB 533* | 0.009 | 0.033 |
| *Mg1655 Δmdtk* | 0.015 | 0.033 |
| *MG1655 RpoB 1304 Δmdtk* | 0.014 | 0.033 |
| *MG1655 gyrB S464Y* | 0.16 | 0.2 |
| *MG1655 gyrB S464Y RpoB 1304* | 0.35 | 0.4 |
| *MG1655 gyrB S464Y RpoB 1279* | 0.35 | 0.4 |
| *MG1655 gyrB S464Y RpoB 1275* | 0.25 | 0.3 |

Ciprofloxacin MIC's and MBC's against strains with various RpoB substitutions: MIC and MBC were tested in two different backgrounds, in MG1655 GyrB S464Y background and in the MG1655. MBC and MIC were determined with approximately $5 \times 10^5$ cell inoculum in 3 ml of media.

**Figure 1:**
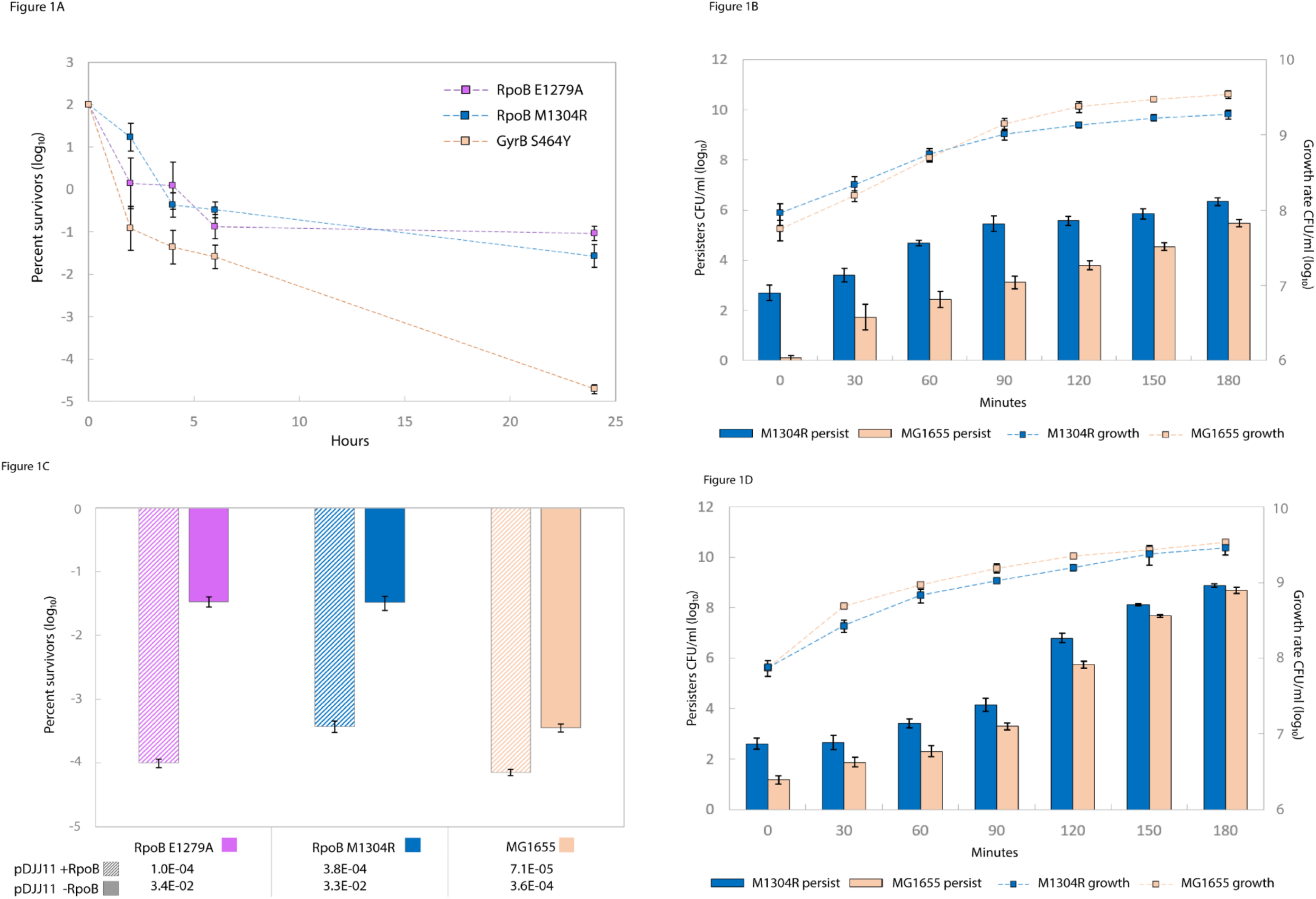
Ciprofloxacin-selected *rpoB* mutations confer high persistence phenotype: (**A**) Survival curves in presence of 4mg/L ciprofloxacin of a parent strain carrying the GyrB S464Y substitution (●) and two RpoB mutants in the same genetic background: RpoB E1279A (●) and RpoB M1304R (●). (**B**) Genetic complementation: RpoB E1279A (●) and RpoB M1304R (●) and GyrB S464Y (●) with a pDJJ11-*rpoB*. (**C**) Genetic complementation: RpoB E1279A (●) and RpoB M1304R (●) and GyrB S464Y (●) with a pDJJ11-*rpoB*. (**D**) Comparison of persistence to ampicillin (18h, 100 mg/L) between MG1655 (●) and RpoB M1304R (●) along the growth curve.

To determine if the high survival frequency phenotype was caused by RpoB M1304R substitution alone or it required a combination with GyrB S464Y, we transduced the *rpoB* mutant allele into a clean MG1655 background and tested the survival frequency of the resulting strain at 10x respective ciprofloxin MIC. Without the *gyrB* mutation, the allele encoding RpoB M1304R increased the survival frequency by a factor of 1,000 (Fig.1B). The wild type *rpoB* expressed from a plasmid fully reversed the phenotype (Fig. 1C). The survival phenotype was not limited to ciprofloxacin. The strain carrying RpoB M1304R substitution yielded 20-25 times as many survivors as its parent strain when treated with ampicillin at 100 mg/L (Fig. 1D). (The RpoB E1279A mutant had a pronounced growth defect and was excluded from further analysis.)

### RpoB M1304R substitution mimics the (p)ppGpp response

Using RNA-seq we identified 379 genes that were differentially expressed (FDR < 0.01) between a strain carrying RpoB M1304R substitution and its parent (Sup. File 1). These genes represented 116 regulons, which were compiled from RegulonDB and EcoCyc (30,31). Of the 116, 12 regulons were represented by at least 10 differentially expressed genes and were significantly enriched (hypergeometric test, adjusted p-value < 0.05) (Fig. 2, panel A). Two regulons stood out from the list of significantly enriched sets. First, the set least likely to be overrepresented by chance was the ppGpp regulon (96 genes out of 167 genes in the regulon were differentially expressed between the two strains, FDR < 0.01). Second, the set with the largest per gene variation in transcript abundances was the DksA-ppGpp regulon. These findings raised the possibility that the effect of RpoB M1304R mimics the effect of (p)ppGpp on transcription and adaptation.

**Figure 2:**
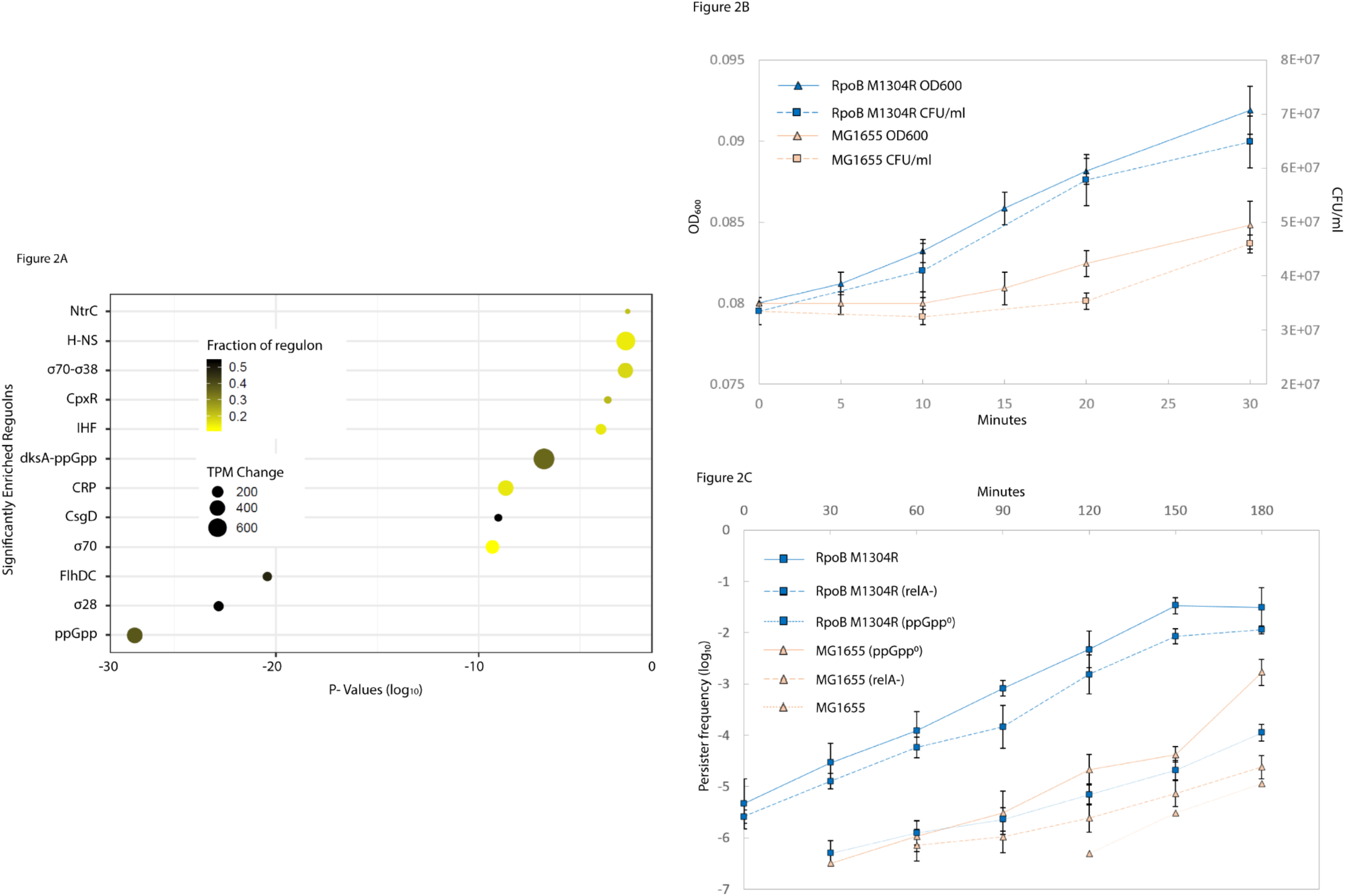
The rpoB M1304R substitution mimics (p)ppGpp response: (**A**) Significantly enriched regulons in RpoB M1304R mutant characterized by the fold change and fraction of the regulon effected by the mutation. (**B**) An adaptation lag of MG1655 parent (●) and MG1655 RpoB M1304R (●) measured in OD_600_ and CFU/ml. (**C**) Ciprofloxacin persistence of RpoBwt(●) and RpoB M1304R (●) in MG1655, *ΔrelA* and *ΔrelA ΔspoT* (ppGpp^0^) backgrounds.

One of the salient features of the ppGpp-dependent response is an adaptive lag during the amino acid downshift (32,33). We compared the lags of the RpoB M1304R strain and its parent MG1655. Upon the downshift, the parent strain exhibited a 20-min delay before resuming growth whereas the mutant started recovering in 5-10 minutes (Fig. 2B). This shortened lag could be due to elevated transcription of amino acid biosynthetic genes in the *rpoB* mutant prior to amino acid downshift. To test this possibility, we examined the activity of *ilvL*::GFP fusion in a flow cytometry experiment. We found that the isoleucine promoter was constitutively up-regulated in the *rpoB* mutant but not in its isogenic parent (Sup. Fig. 1), strongly suggesting that the C-terminal RpoB mutation “behaves” like a typical stringent mutation, which is usually found between 526-533 aa residues of RpoB.

However, since the stringent-like phenotype in the M1304R strain was observed in ppGpp+ cells, it was also possible that the mutant has a higher basal level of ppGpp. We tested this possibility by directly measuring ppGpp levels in cell extracts. The alarmone abundance was indistinguishable between the parent and the *rpoB* mutant in presence of amino acids (Fig. 3A). After 1h amino acid downshift, ppGpp levels increased 10 times over the basal level in both strains (Fig. 3B). Thus, the RpoB M1304R substitution either makes RNA polymerase more sensitive to ppGpp or mimics the response in a ppGpp-independent manner.

**Figure 3:**
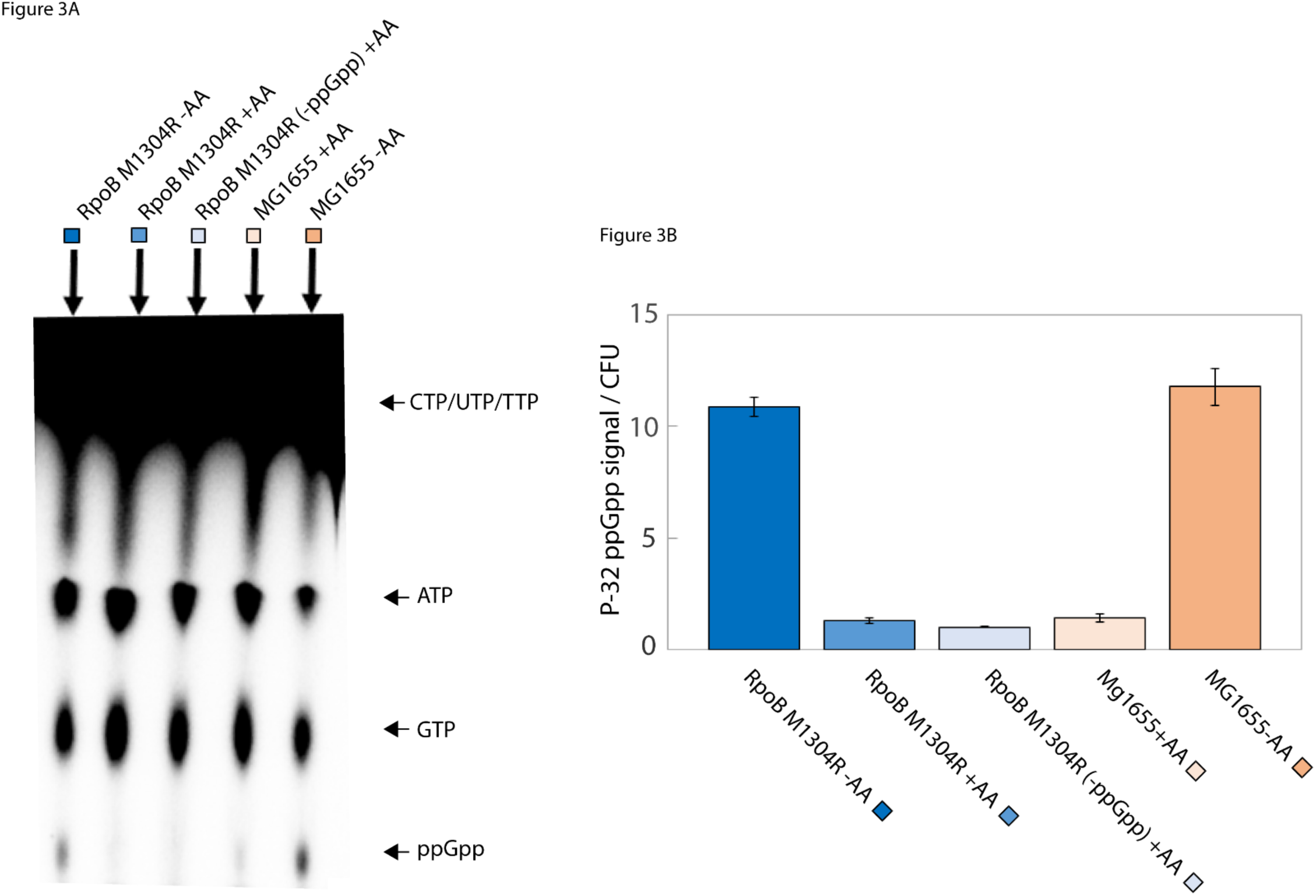
ppGpp mimicry is not a result of ppGpp accumulation: Prior to the experiment all strains were grown in the presence on amino acids, at the start of the experiment cells were washed and placed in the media containing ^32^P with (+AA) or lacking (-AA) amino acids. (**A**) Radiolabeled ^32^P nucleotide TLC plate comparing MG1655 RpoB M1304R +/- amino acids, to MG1655 RpoB M1304R ppGpp^0^ + amino acids, to MG1655 +/- amino acids. A faint, ^32^P trace in the middle three lanes is the background noise (34). A small accumulation of ppGpp can be observed in the fourth lane as a result of cultures’ entry into early stationary state. (**B**) Fiji analyzed TLC plate and normalized to corresponding CFU/ml microbial counts: MG1655 RpoB M1304R (●) -AA, MG1655 RpoB M1304R (●) +AA, MG1655 RpoB M1304R ppGpp^0^ (●), MG1655 +AA (●), MG1655 –AA (●).

We tested the ppGpp dependence of RpoB M1304R by transducing the mutation into the *relA*- and *relA*-*spoT*-(ppGpp^0^) backgrounds. While the removal of *relA* alone slightly reduced the frequency of survivors compared to the RpoB M1304R *relA*+ parent, the RpoB M1304R persistence phenotype without ppGpp was similar to that of the wild type (Fig. 2C). Thus, the *rpoB* persistence mutation, while being dependent on ppGpp for its full effect, makes the basal persistence independent of the alarmone. Such independence, or ppGpp mimicking, can be manifested by phenotypic rescuing of the amino acid auxotrophy of ppGpp^0^ mutants (19, 35). However, we found that the RpoB M1304R ppGpp^0^ mutant still required amino acids for growth (data not shown).

### Bone fide rpoB stringent mutations cause high-persistence (hip) phenotype

To determine if stringent *rpoB* mutations (27), which are known to rescue the ppGpp^0^ auxotrophy, can also rescue the persistence phenotype, we selected rifampicin resistant, *rpoB* stringent mutants in a ppGpp^0^ strain (“Materials and Methods”) and characterized persistence of one of them, RpoB L533P. While ppGpp^0^ yielded no persisters between OD_600_ 0.3 −1.6, RpoB L533P ppGpp^0^ had as many persisters throughout the growth curve as wild type MG1655 in the early stationary phase (Fig. 4A and B).

**Figure 4:**
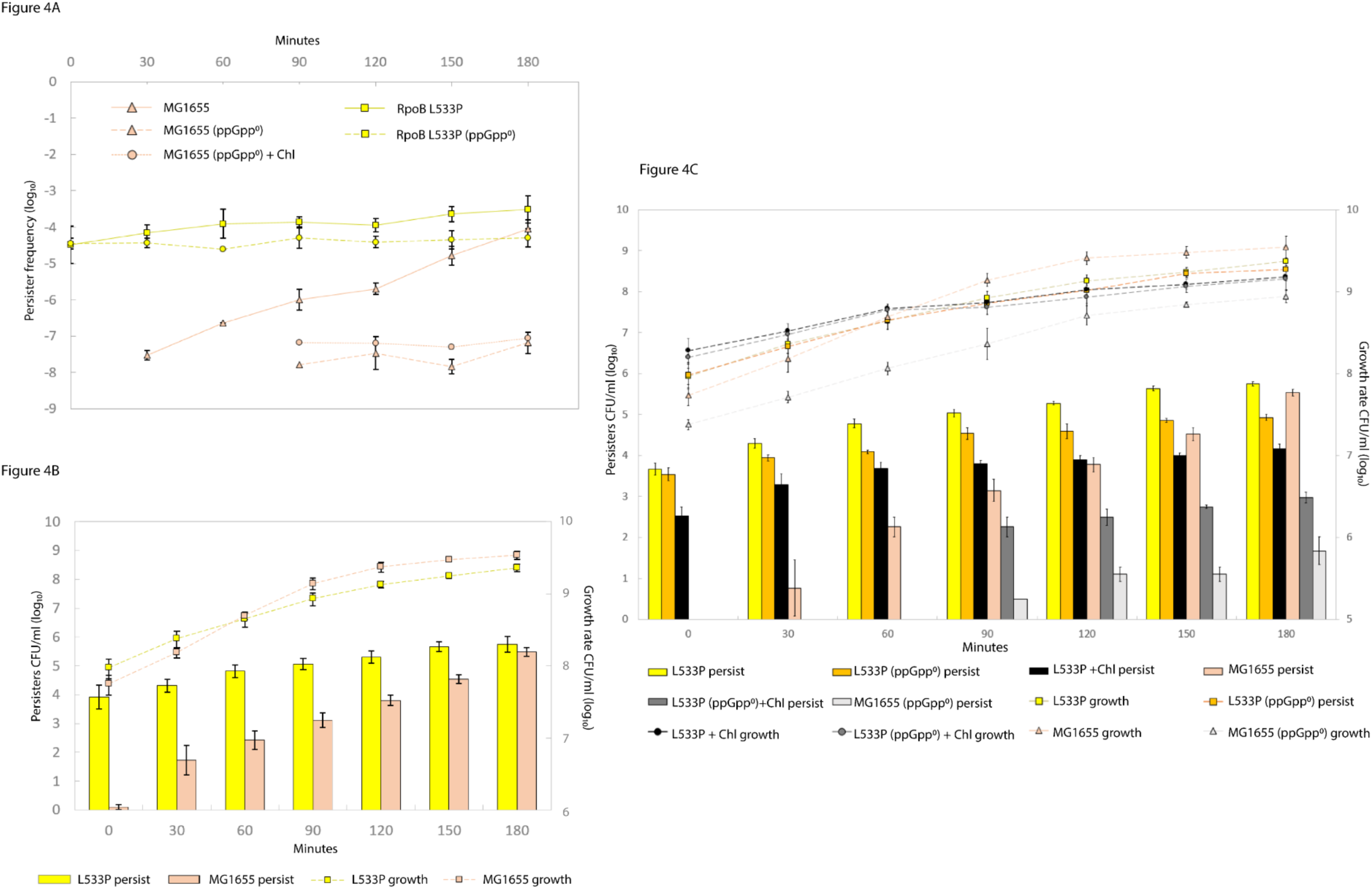
*rpoB* stringent mutations confer high persistence phenotype that can’t be explained by a slower growth: (**A**) Persister frequency comparison between the MG1655 RpoB L533P (●), MG1655 RpoB L533P ppGpp^0^ (●), MG1655 (●), MG1655 ppGpp^0^ (●) as well as MG1655 ppGpp^0^ (●) slowed with 0.63 mg/L chloramphenicol. The persistence phenotype of stringent mutant is consistently maintained at 1:10,000 −50,000 and is independent of ppGpp.(**B**) Comparison of persistence to ciprofloxacin (18h, 1mg/L) between MG1655 (●) and MG1655 RpoB L533P (●) along the growth curve. Persistence exhibited by stringent mutant is independent of the growth state. **(C)** Comparison of persister formation between six different strains throughout the exponential and stationary states, in the presence and absence of chloramphenicol (Chl). MG1655 RpoB L533P (●), MG1655 RpoB L533P (ppGpp^0^) (●), MG1655 (●), MG1655 RpoB L533P +Chl (●), MG1655 RpoB L533P (ppGpp^0^)+Chl (●), MG1655 (ppGpp^0^) (●).

Interpretation of the persistence data from strains carrying RpoB M1304R and RpoB L533P substitutions is confounded by a relatively slower growth of both mutants (Table 2). To evaluate the effect of growth rate on persistence, we measured the frequency of persisters in MG1655 at OD_600_ of 0.5 slowed by chloramphenicol to a 22, 27, 30, 35- and 40-min doubling time. Predictably, there was an inverse relationship between growth rate and persister frequency. Yet, when MG1655 was slowed down to the RpoB L533P level, the stringent mutant still yielded over 600 times as many persisters (Sup. Fig. 2). The higher persistence observed in slow-growing cultures could also be a consequence of the ppGpp effect, which controls the growth rate (36), and not of the slow growth itself. To test this, the growth of the ppGpp^0^ parent was halved by chloramphenicol (Sup. Fig. 3A) and the survival frequency was measured at different points along the growth curve. The slowing down had no measurable effect on persistence (Fig. 4A), implying that ppGpp is required for the effect of growth rate on persistence. This straightforward conclusion, however, may not be possible if chloramphenicol kills a substantial fraction of cells and the persistence is measured only over the surviving sub-population. We tested whether this is the case by plating MG1655, RpoB1304 and RpoB533 (both in MG1655 background) as well as their ppGpp^0^ derivatives on plates containing 0, 0.5, 1, 2, 4 and 25 mg/L of chloramphenicol, as was done in (37). We found that chloramphenicol had no effect on CFU’s at concentrations between 0 and 4 mg/L (Sup. Fig. 3B and C) and hence the aforementioned conclusion stands.

**Table 2:** Doubling times.

|  | Doubling time |
| --- | --- |
| Strains | Minutes |
| *MG1655* | 20±1.2 |
| *MG1655 RpoB L533P* | 27.2±2.3 |
| *MG1655 RpoB M1304R* | 25.2±1.4 |

Growth rates of strains with RpoB substitutions in MG1655 background: Doubling time was determined by CFU/ml.

We next asked what effect would slowing down high persister mutants by sub-inhibitory concentrations of chloramphenicol would have on the frequency of persisters? We expected to see an even higher levels of persistence in RpoB L533P and RpoB L533P(ppGpp^0^) mutants in presence of the translation inhibitor. Surprisingly, the opposite was the case. Persistence in the RpoB L533P mutant had decreased by almost two logs (93-97% depending on the time point). The drop in the persistence was even greater when it came to the RpoB L533P ppGpp^0^ mutant: no observable persistence in the first 90 minutes and over two logs decrease in persistence over the remaining time points (Fig. 4C).

### Indole contributes to the RpoB-mediated persistence

To determine if there are any commonalities between the molecular phenotypes of strains carrying RpoB L533P and RpoB M1304R mutations, we compared their transcriptional profiles in wild type and ppGpp^0^ backgrounds. Among 697 significantly differentially expressed genes, one operon, *tnaCBA*, stood out as the most upregulated (Sup. File 2). We confirmed activation of the *tnaC* promoter in both of the *rpoB* mutants by flow cytometry (Fig. 5A). Of note, the distribution of the fluorescent signal in the mutants was quantitatively and qualitatively different from that in the parent: not only it was shifted to the right, but it also had two modes, with the second mode representing about 3-5% of all fluorescent counts with about 10-times higher fluorescence than the corresponding mutant average. A *tnaA* knockout in RpoB L533P and RpoB M1304R mutants reduced the frequency of persisters by 50% and 80%, respectively (Fig. 5B). Concentration of indole in the RpoB L533P and RpoB M1304R cultures was 15 μM and 50 μM, respectively, compared to undetectable levels in the parent. However, when we added up to 50 μM indole to a wild-type culture we did not observe any increase in persister formation. When we added indole to a final concentration of 500 μM (a stationary culture equivalent), persistence increased 7-10 times. Given the distribution of the activity of *tnaC* promoter in a population of the *rpoB* mutant cells, it is plausible that only 3-5% of the population produce indole in quantities sufficient to trigger the dormant state, which in turn will happen only in a fraction of that sub-population.

**Figure 5:**
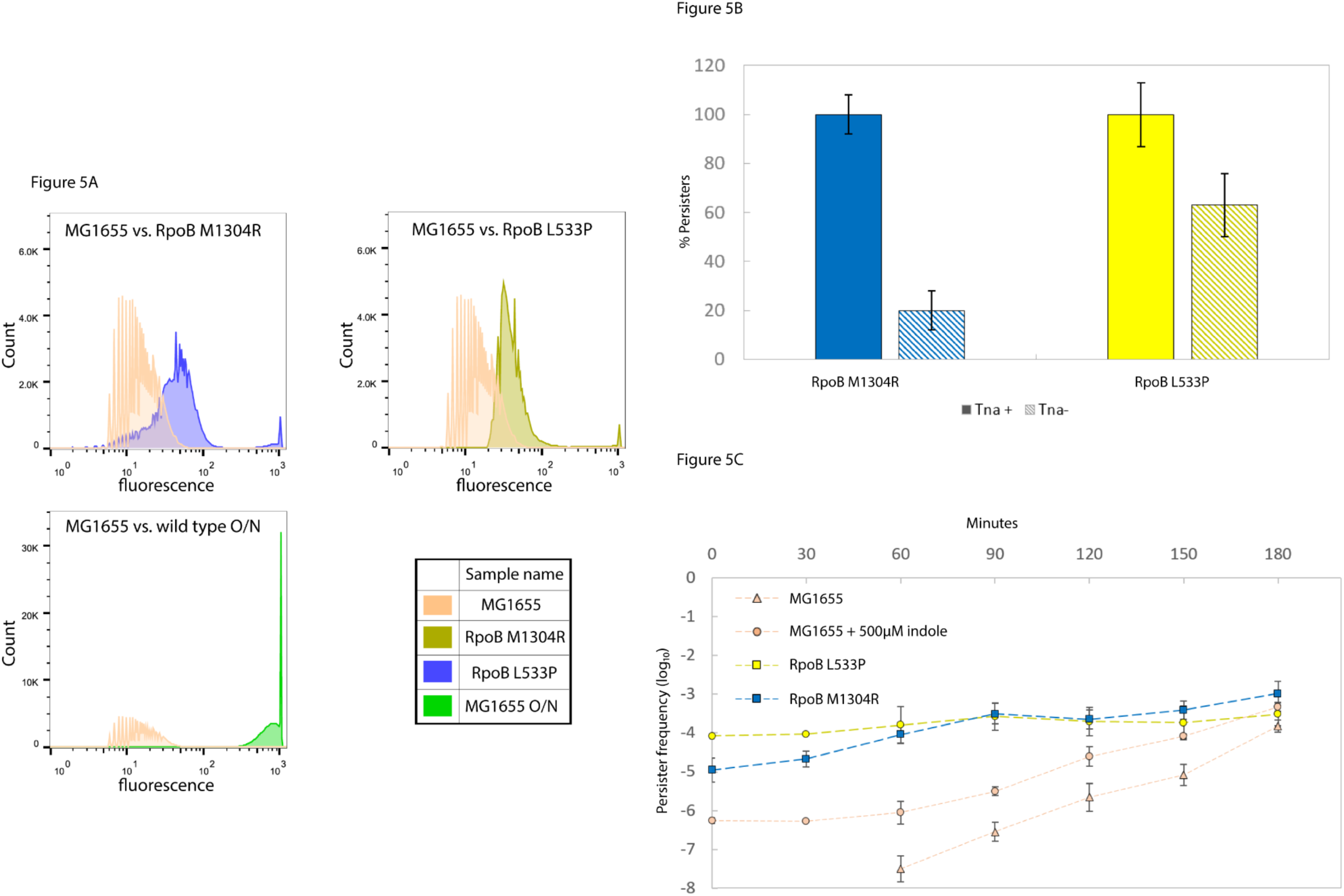
The effects of indole on persistence: (**A**) Activation of *tnaC(p)* in MG1655 RpoB M1304R (●) and MG1655 RpoB L533P (●) compared to MG1655(●). The bottom left panel compares the activity of the *tnaC* promoter between exponential and overnight MG1655 cells (●). (**B**) Effect of *tnaA* knockout on ciprofloxacin persistence in MG1655 RpoB M1304R (●) and MG1655 RpoB L533P (●) was measured by averaging percent survival of *tnaA* across 7 time points (from t0 at OD_600_ of 0.3 all the way to stationary state). At each time point fraction of survivors was determined for *rpoB* mutant and its *tnaA* null counterpart. The error bars represent standard deviation of survivor fraction across all seven time points (**C**) Ciprofloxacin persistence of MG1655 RpoB M1304R (●), MG1655 RpoB L533P (●), MG1655 (●) and MG1655 + 500μM (●).

### RpoB high-persistence mutations are found among clinical isolates

We used sequences stored in PATRIC database (38) to determine if the *rpoB* high-persistence mutations occur in clinical isolates. Since in our selection mutations in *rpoB* followed a *gyrB* mutation, we first analyzed the co-occurrence of *gyrB* mutations known to contribute to quinolone resistance (39,40) and *rpoB* mutations occurring at or in the vicinity of 1279 and 1304 residues (41). Sequences were processed and the statistical significance of co-occurrences was analyzed as described in (42). Although we did not find E1279A and M1304R among RpoB sequences of *E. coli* clinical isolates, W1275G and T1286P substitutions did significantly co-occur with D426N and S464Y substitutions in GyrB (Chi-squared test, adjusted p-value = 0.002). We introduced the mutation resulting in W1275G substitution into the *E. coli* chromosome. RpoB W1275G mutant yielded nearly as many persisters as the M1304R strains (Sup. Fig. 4). (We were unable to construct a strain with RpoB1286P substitution.)

Although rifampicin is not normally used against *E. coli* infections, *rpoB* sequences from 22 clinical isolates carried known Rif^R^ mutations (43). Three out of the 22 (13.6%) had substitutions in Ser531 residue, which is a known stringent mutation (35) and exhibited high persistence (similar to L533 substitution) when transferred into MG1655 (data not shown). Coincidently, all three mutations were found in the isolates that also carried known *gyrA* (40) and/or *gyrB* (44) quinolone resistance mutations.

Next, we investigated if stringent mutants could outcompete other Rif^R^ mutants in presence of bactericidal antibiotics. We selected 104 Rif^R^ mutants in MG1655 ppGpp^0^ background, so that the stringency could be easily confirmed by plating on amino acid-less M9 plates. Out of 104 rifampicin resistant mutants, only 3 were stringent. After pooled colonies were treated with ciprofloxacin, the fraction of stringent mutants increased to 23%, compared to the 3% observed after the rifampicin selection, suggesting that stringent mutations can confer competitive survival advantage.

We also determined the frequency of stringent mutations among the Rif^R^ isolates of pathogens that are routinely treated by rifampicin, *A. baumannii, M. tuberculosis*, and *S. aureus*. The prevalence of the mutations that are homologous to *E. coli* Rif^R^ stringent was even higher than what we had observed in *E. coli*: 96.2%, 23.3%, and 50.9%, respectively. To determine if such mutations can confer high persistence, we investigated the effect of the RpoB S586L substitution in MRSA (an equivalent of S531L substitution in *E. coli*). Stringent mutants were selected as described in “Materials and Methods”. The persister frequency of the RpoB S586L mutant was more than 10,000 times higher than that of the parent in the early exponential phase, and even in the late stationary phase the mutant produced 8 times as many persisters (Fig. 6A, B).

**Figure 6:**
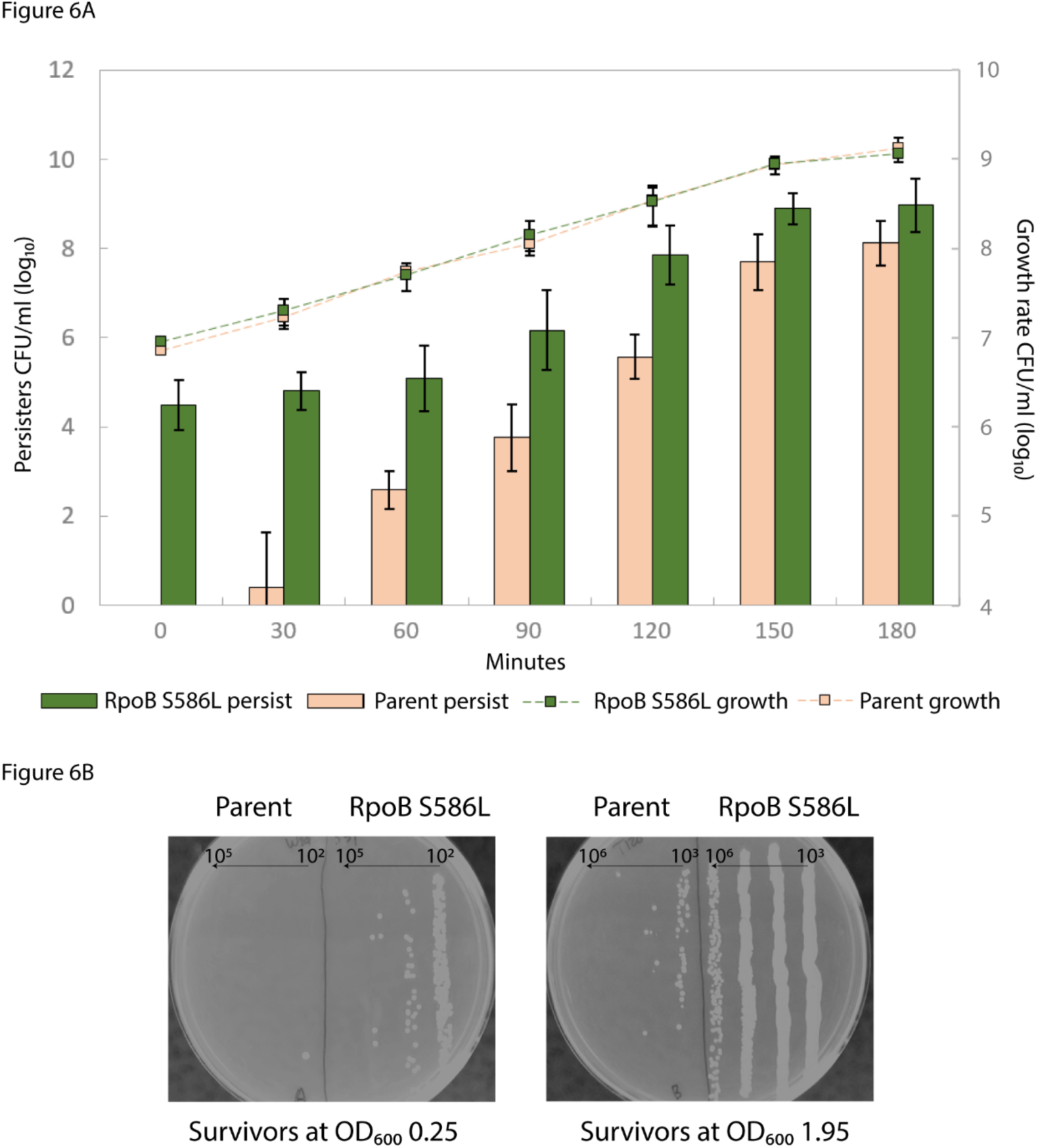
An *rpoB* stringent mutation confers high persistence in *Staphylococcus aureus*: (**A**) Comparison of ampicillin persistence (18h, 100 mg/L) between MRSA WCUH29 (●) and WCUH29 RpoB S586L (●). (**B**) Photographs of plates with survivors during exponential phase (*left*) and early stationary phase (*right*), the left side of each plate is the parent, and the right is the *rpoB* stringent mutant. Dilution ranges are shown above the horizontal arrows on each plate.

## Discussion

Bacteria adapt to antibiotic-containing environments by acquiring resistance mutations. Here, we demonstrated that some of these mutations, including those that are highly prevalent in clinical isolates, can confer an additional benefit of increased persistence to bactericidal antibiotics. These high-persistence mutations resided in two regions of the beta subunit of the RNA polymerase: 1275-1304 and 531-533 and 526 *E. coli* equivalent in MRSA. Mutations in the first region can be readily selected at low concentrations of ciprofloxacin and those in the second are commonly occurring rifampicin-resistance mutations. Thus *hip* mutations can be fixed in a bacterial population not only through cyclical exposure to bactericidal antibiotics (45,46,47), but also as a result of pleiotropy of certain antibiotic resistance mutations.

The RpoB M1304R substitution resulted in two key phenotypes: reduced susceptibility to ciprofloxacin by 1.5-2-fold and increased persistence by up to a factor of 1,000. The two effects depend on different mechanisms, however. Among top 50 significantly over-expressed genes as a consequence of RpoB M1304R substitution, the function of only one of them, *mdtK*, which encodes a multi-antimicrobial extrusion transporter, could be directly linked to drug resistance (48) (Sup. File 3). Pietsch *et al*. found that similar *rpoB* mutations increased fluoroquinolone MIC by upregulating *mdtK* (41). However, while an *mdtK* knockout in RpoB M1304R strain resulted in the full reversal of the resistance phenotype (Table 1), it had no effect on persistence (Sup. Fig. 5), suggesting that the two phenotypes can be segregated.

MG1655 strain carrying RpoB L533P substitution, which confers rifampicin resistance, had no measurable effect on ciprofloxacin MIC. And yet it resulted in a striking persister phenotype; the frequency of persisters was independent of the growth phase (Fig. 4A).

Another relevant phenotype exhibited by the mutants is a slower growth. Decreasing the growth rate resulted in an increased number of persisters. However, the persister frequency of the *rpoB* rifampicin resistant mutant was more than 600 times higher when persistence was measured using ciprofloxacin and more than 80 times higher when using ampicillin, compared to what would be expected from the reduced growth rate alone in both ppGpp+ and null cells (Fig. 4A, Sup. Fig. 2). These findings strongly argue against the notion that a decreased target abundance, which should be a consequence of slowed growth, is sufficient to explain increase in persistence or that it confounds the phenomenon altogether.

The *rpoB* mutations completely overrode the ppGpp requirement for persister formation. This strongly suggests that the persister state is the result of a transcriptional program and ppGpp effects persistence primarily, if not solely, as a modulator of transcription. The mutants also illuminated the importance of differences in transcriptional activity among individual cells as an attribute of the *hip* phenotype: a sub-population of mutant cells had the promoter of the *tnaA* gene upregulated at least 10 times, while most of the population showed only marginal shift in the promoter activity. We demonstrated that this *tnaA* activation resulted in accumulation of indole that accounted for 50-80% of the *hip* phenotype in the *rpoB* mutants. Although these results are qualitatively consistent with earlier observations of the effect of indole on persistence (5, 6), we also found that indole had no effect on persistence without ppGpp or its transcriptional mimics. The role of *tnaA* and indole in the persiter state formation may tie together the metabolic state of individual cells with transition to dormancy. The low-affinity tryptophan transporter TnaA is activated when tryptophan is used as a carbon source (49,50), which happens only when all other preferred carbon sources in rich media have been exhausted (51,52). Switching to tryptophan, or other “lesser” carbon resources, may signal imminent starvation which, in combination with other yet unknown factors, establishes the persister state. In the *rpoB* mutants, *tnaA* transcription is constitutively upregulated in a number of cells, resulting in the “out-of-order” tryptophan utilization, starvation signaling by means of indole, and dormancy. This explanation also applies to indole-free species, where the “quality” of carbon and nitrogen source facilitates state transition. For example, Halsey *et al*. have suggested that histidine catabolism may be involved in regulation of CcpA response in *MRSA* (53,54,55), a major transcriptional regulator involved in metabolic regulation under suboptimal growth conditions, virulence, antibiotic resistance and biofilm formation (56,57).

The primary effect of rifampicin resistance mutations is to provide protection against the drug. Over one hundred mutations in RNA polymerase have been shown to confer varying degrees of rifampicin resistance (58). However, 17 mutations, which map between 505-690 amino acid positions of RNA polymerase β-subunit, make up the majority of the clinically relevant variants (59). About half of these mutations appear to have no growth defect (25). Yet, growth-defective stringent mutants can be frequently found in pathogenic species (60,61), indicating that stringent mutations must be providing sufficient adaptive/survival advantage even before the mutants have a chance to accumulate compensatory mutations restoring their growth fitness (62). Our findings suggest that persistence may be one of such advantages.

Finally, rifampicin is almost always used in combinations with other bactericidal antibiotics against some of the more antibiotic-recalcitrant bacteria (63). From adaptation prospective, the most direct path to surviving an onslaught by several antibiotics may go through acquisition of a single *hip* mutation rather than multiple resistance mutations to different antibiotics. Hence, selecting for *rpoB*-dependent persisters may promote the establishment of chronic infections (64), rather than subverting them, in a long run. However, there may be a simple and elegant solution to revering the mutations’ effect. The drastic decrease in persistence which we observed in the MG1655 RpoB L533P and its ppGpp^0^ derivative treated with sub-inhibitory chloramphenicol concentrations suggests that transcriptional dysregulation induced by *rpoB* mutations may be counteracted at a translational level. Just as with any phenotypic switch, expression of genes contributing to induction of the persister state has to reach a certain threshold. Compromising the cell’s ability to reach that threshold may negate the phenotype.

## Materials and Methods

### Bacterial strains and culture conditions

Antibiotics, strains, and plasmids used in the study are listed in Supplementary Tables 1-3. *E. coli* strains were derived from K-12 MG1655 (65) by selection, P1 transduction or λ-red recombination (66,67). *E. coli* cultures were grown in LB or MOPS-glucose (33), *S. aureus* – in tryptic soy broth/agar (TSB/TSA) (68) at 37°C with aeration at 250 rpm.

### Determining MIC and MBC

MIC (Minimum Inhibitory Concentration) and MBC (Minimum Bactericidal Concentration) estimates were obtained according to standard protocols (69). MIC was determined by inoculating 5 x 10^5^ (± 2 × 10^5^) cells into 3ml Mueller-Hinton containing a range of drug concentrations (1, 0.5, 0.25, 0.125, 0.062, 0.031, 0.015, 0.008, 0.004 mg/L) and checking turbidity the following day. After establishing initial MIC range, the procedure was repeated until MIC was determined within 0.001 mg/L for the mutants in the MG1655 background, and within 0.02 mg/L for mutants in the MG1655 GyrB S464Y background. To measure MBC, the same cultures that were used for MIC measurements were washed and plated for survivors. Prior to addition of the antibiotic all cultures were plated by serial dilution to determine the starting CFU/ml count. The concentration corresponding to a 3-log drop in viability was recorded as the MBC for the corresponding strain. The MICs and MBCs for each strain were obtained from testing at least 3 independent biological samples. Both MIC and MBC measurement were also conducted in LB media (70), resulting in identical values across all mutants.

### Persistence measurement

Persistence was measured following a protocol described in (71). Bacteria were maintained by repeated dilutions in an early exponential phase between an OD_600_ 0.05-0.3 for at least 10 generations. Upon reaching an OD_600_ of 0.3±0.03 after the 10^th^ doubling, a 3ml sample was taken out every 30 min for the next 3 h and ciprofloxacin or ampicillin was added to a final concentration of 1 mg/L (or 4 mg/L in the case of *gyrB* mutants to assure killing at 10X MIC) and 100 mg/L, respectively. After 17h incubation, samples were plated for CFU. Persistence was determined as a number of survivors per ml normalized to the CFU/ml counts of the culture at the time of antibiotic application.

### ppGpp quantification using TLC

Overnight cultures were diluted 1:1,000 in 3.8ml of glucose-MOPS supplemented with 20 L-amino acids and grown to an OD_600_ of 0.3. Cultures were then split into two 1.5ml tubes and spun down. Half of the sample was resuspended in fresh 3ml MOPS supplemented with amino acids, and another half was used to inoculate amino-acid-free MOPS to a starting OD_600_ of 0.15 for all samples. From each sample CFU/ml were determined by serial dilution. All samples were then supplemented with 20μCi of ^32^P orthophosphate. Following 1h incubation cells were lysed and 20 μl of lysate was spotted on a nitrocellulose TLC plate. Samples were mobilized by pH3.5 water-based phosphate buffer (72). Samples were visualized by Amersham Typhoon control software and analyzed using Fiji software (73).

### Single gene expression assay and flow cytometry

Strains were transformed with a pUA66 plasmid carrying a *ilvl-p*:*gfp, hisL-p:gfp, tnaC-p:gfp* and *mdtK-p*:*gfp* promoter fusion (74). Overnight cultures of newly-obtained transformants were diluted 1:1,000 into pre-warmed glucose-MOPS supplemented with 20 L-amino acids, with each amino acid present at a final concentration of 0.04 mg/L, and maintained for at least 10 generations in an early exponential phase as described above. When an OD_600_ of 0.3 was reached after the 10^th^ doubling, cultures were diluted 100 times for flow cytometry. The GFP signal was measured at 14-66 μl/min flow rate. Data were analyzed using FlowJo. Fluorescence was normalized to both OD_600_ and CFU/ml. The *tnaC-p:gfp* experiment was carried out in LB and GFP signal was measured in saline.

### Whole Genome Sequencing

Isolates were grown overnight from a single colony in LB at standard conditions. DNA was extracted using a Promega Wizard Genomic DNA Purification Kit. Libraries were prepared using a TruSeq Nano library preparation kit. Sequencing was conducted using Illumina HiSeq2500. De-multiplexing was performed using FASTX-Toolkit (http://hannonlab.cshl.edu/fastx_toolkit/index.html). The reads were mapped to the reference genome (GCF_000005845.2_ASM584v2) using Bowtie 2 (78,79). The SAM files of mapped reads were converted to BAM format and sorted using SAMTOOLS (69). Mutation were identified using Breseq (80) and confirmed using standard Sanger sequencing, using primers listed in Supplementary Table 4.

### RNA-seq experiment and analysis

Overnight cultures were diluted 1:1,000 into 50ml LB and maintained in exponential phase as described above. After at least 10 generations in exponential phase and upon reaching an OD_600_ of 0.3, 10ml samples were transferred into 15ml conical tubes containing 1.25 ml of the “stop” solution (ethanol v/v 95%, and phenol v/v 5%). Cells were then spun down and decanted. Pellets were flash frozen in liquid nitrogen and kept at −80°C until all samples were collected. Pellets were thawed on ice and resuspended in 200μl of TE buffer containing 20,000 mg/L of lysozyme and incubated at 37°C for 10 min. Total RNA was purified following a standard RNAeasy mini prep kit protocol (Qiagen, Hilden, Germany).

Before constructing the sequencing library rRNA was removed. The library was made using TruSeq Stranded mRNA library prep kit. Sequencing was done using Illumina HiSeq2500. De-multiplexing was performed using Illumina’s bcl2fastq software 2.20. The reads were mapped to the reference genome (GCF_000005845.2_ASM584v2) using HISAT2 (75). The final per feature read counts were obtained using Rsubread (76). Analysis of differential expression was done using DESeq2-based (77).

### Indole assay

Indole concentration was measured by adding 400 μL of Kovack’s reagent to a 3ml culture of exponential cells at an OD_600_ of 0.3 ± 0.05. Cells were incubated with the reagent for 30 min at room temperature in the dark. 200 μL from the surface alcohol layer was moved into a flat-bottom, 96-well plate and absorbance was measured at 535 nm using a plate reader. A standard curve was constructed using Kovack’s reagent colorimetric assay with following concentrations of indole: 10, 25, 50, 100, 200, 400, 500 μM in LB.

## Acknowledgements

We would like to thank Dr. Ding Jin for pDJJ11 plasmid and Thu Tran for help with initial experiments.

---

**Supplementary figure 1.**
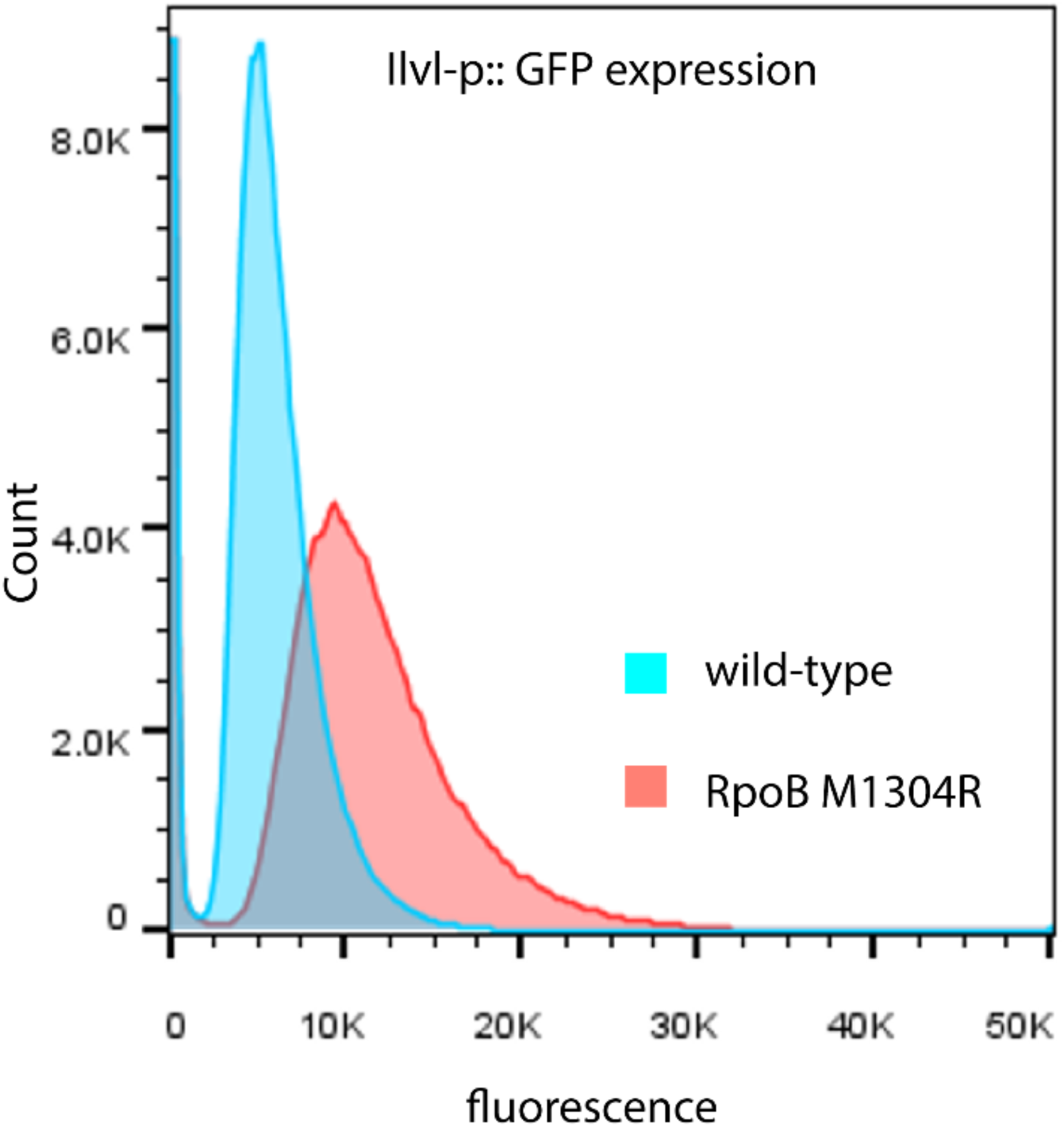
Flow cytometry of ilvL(p)::GFP reporter: cells grown in MOPS medium supplemented with all 0.2% glucose and all 20 amino acids. Samples for flow cytometry were taken after culture had reached OD_600_ of 0.3 and diluted down using the same pre-warmed media to 0.03±0.01. Comparison is between MG1655 wild type (●) and RpoB M1304R (●) in the same backgrounds.

**Supplementary figure 2.**
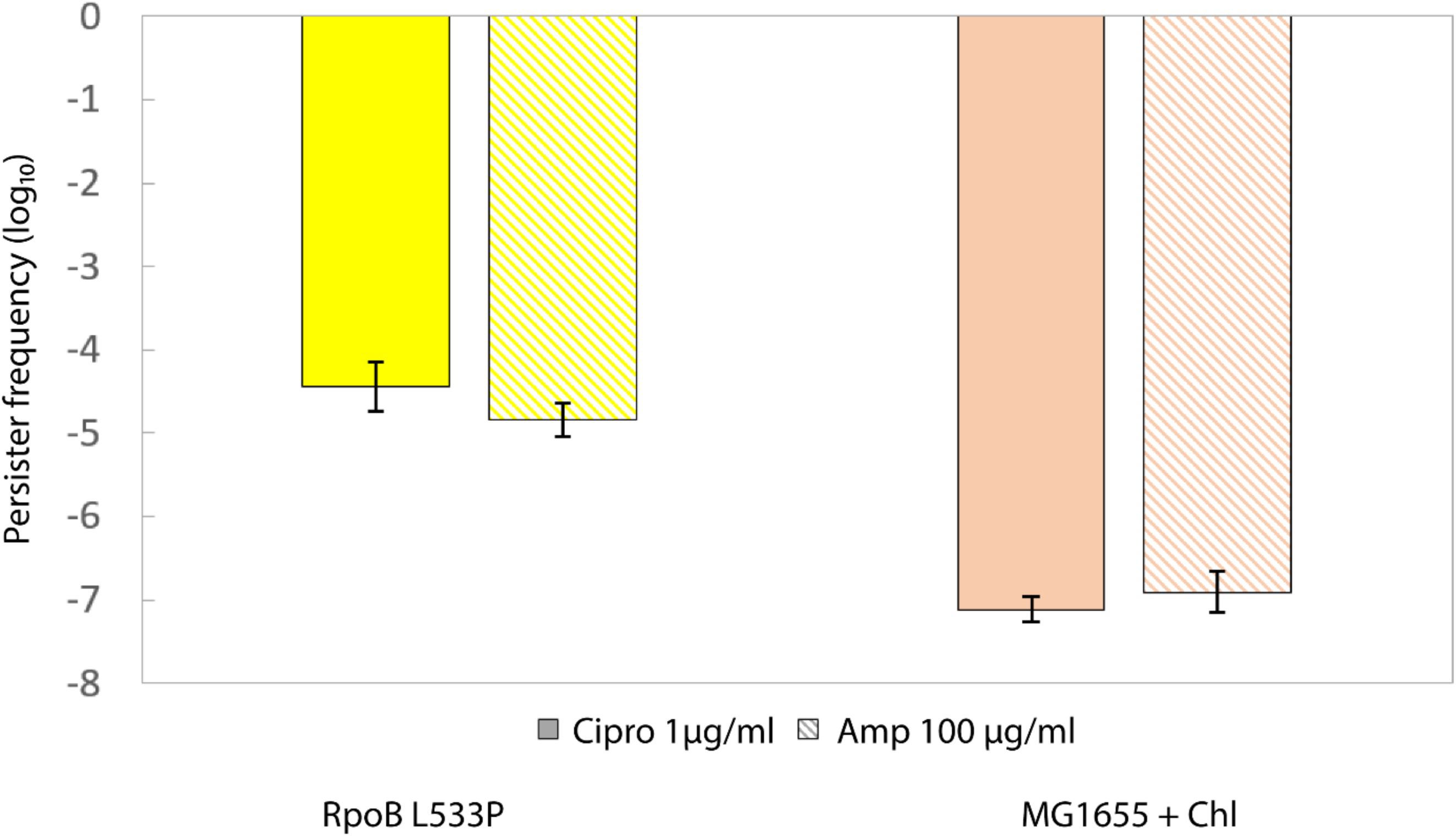
Stringent mutant persistence is not a consequence of slowed growth: Comparison of persistence between MG1655 RpoB L533P (●) and MG1655 (●) slowed to 30 min doubling time using chloramphenicol. The left bar represents persister frequency when strains were treated by ciprofloxacin, and the right bar shows persister frequency following ampicillin treatment. All treatments were performed on exponentially growing cultures at OD_600_ of 0.5.

**Supplementary figure 3.**
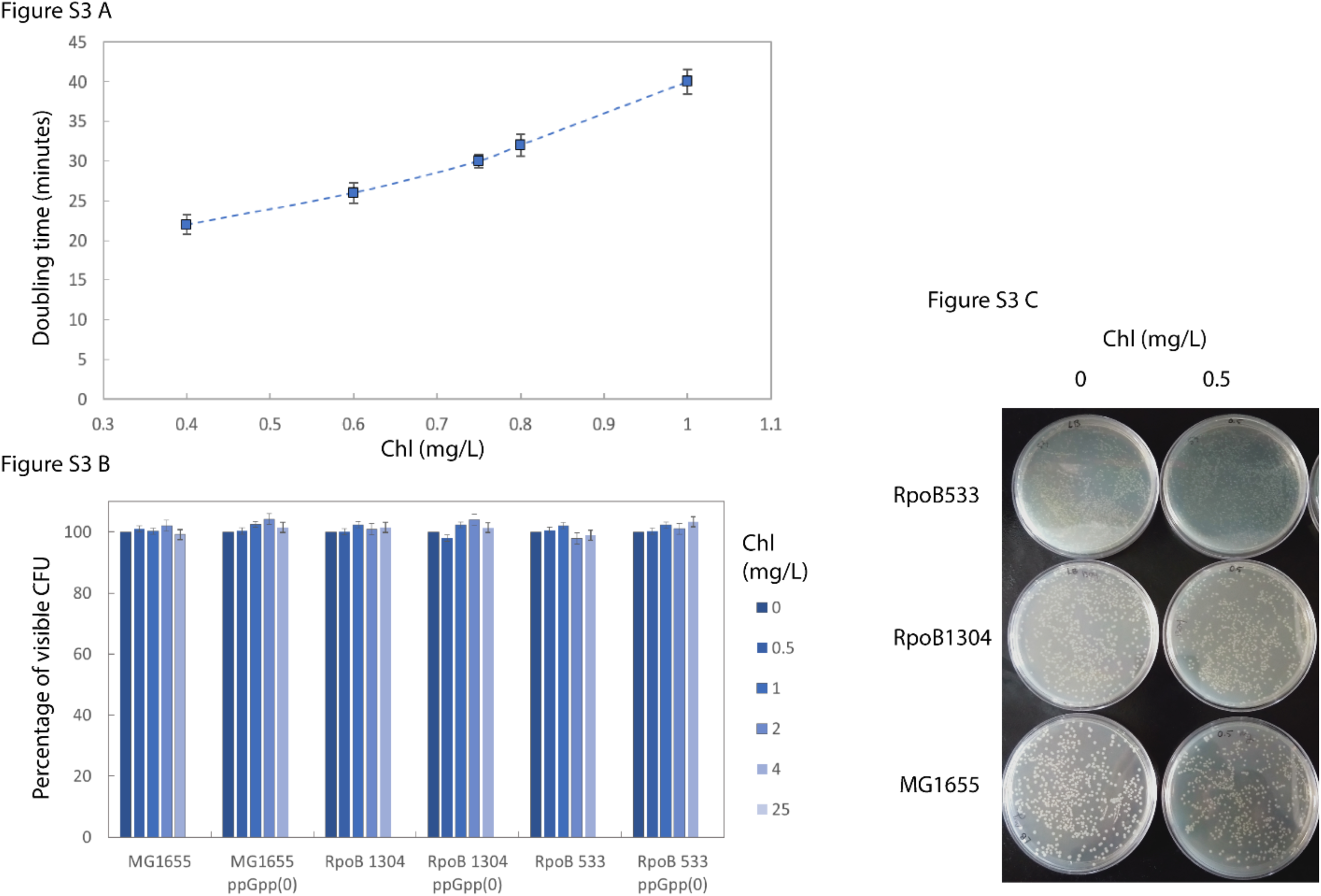
Effects of sub-inhibitory concentrations of chloramphenicol on growth of E. coli: (**A**) MG1655 was grown in presence of 0.4, 0.6, 0.75, 0.8 and 1μg/mL of chloramphenicol and the doubling time at each concentration was determined using CFU/ml. (**B**) Percentage of visible CFU counts plated on LB plates containing sub inhibitory concentrations of chloramphenicol (0, 0.5, 1, 2, 4, 25 mg/L) for the six strains used in the study. All strains tested were in MG1655 background. Error bars represent standard deviation from three biological replicates (**C**) A photograph depicting MG1655, RpoB1304 and RpoB533 both in MG1655 background plated on LB plates containing 0 and 0.5 mg/L of chloramphenicol.

**Supplementary figure 4.**
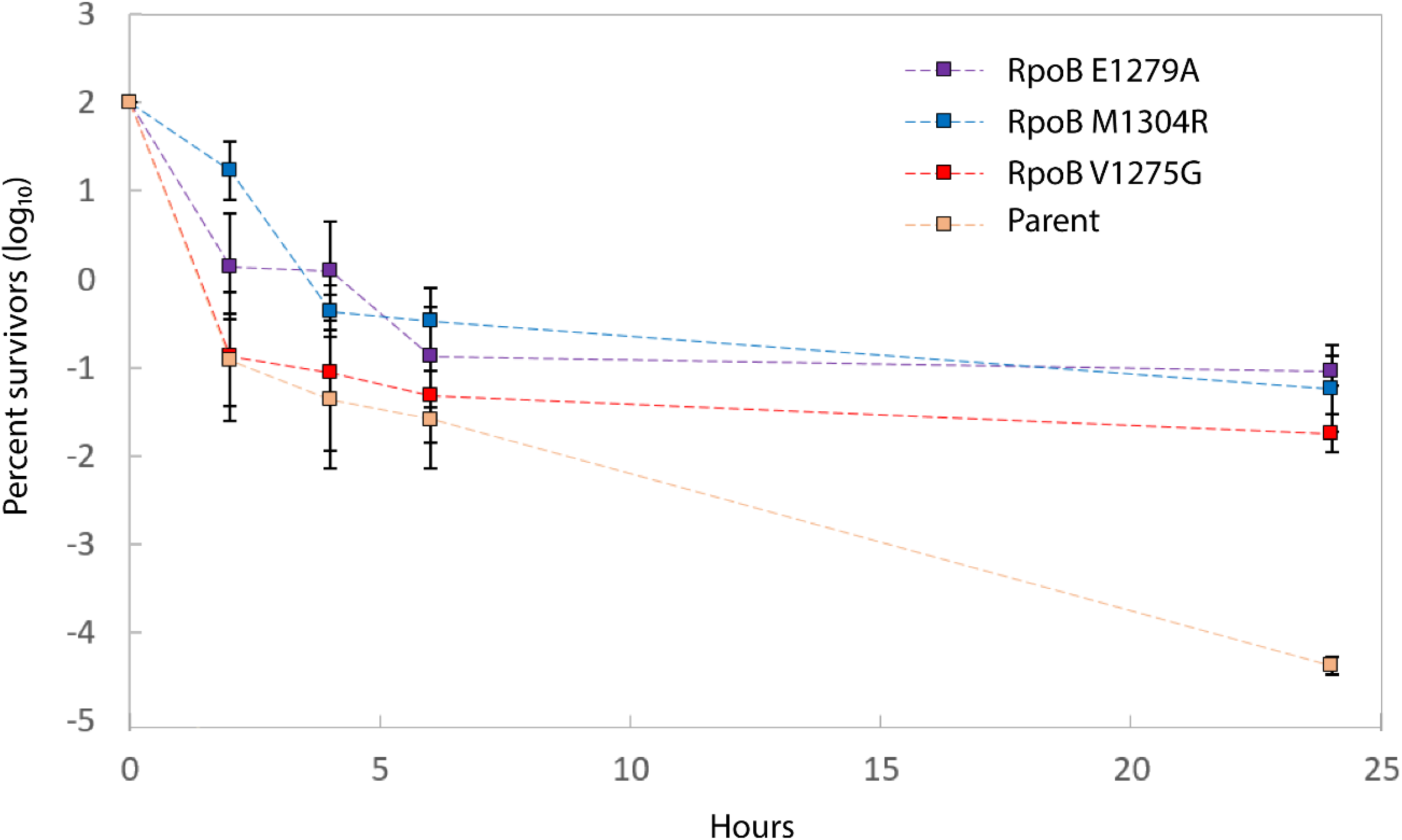
The *rpoB* mutations co-occurring with gyrase mutations from clinical isolates confer increased persistence. Ciprofloxacin (4μg/mL) 24h kill curves comparing the parent strain with a GyrB S464Y substitution (●) to the mutants in the same background carrying E1279A (●), M1304R (●) and V1275G (●) RpoB substitutions.

**Supplementary figure 5.**
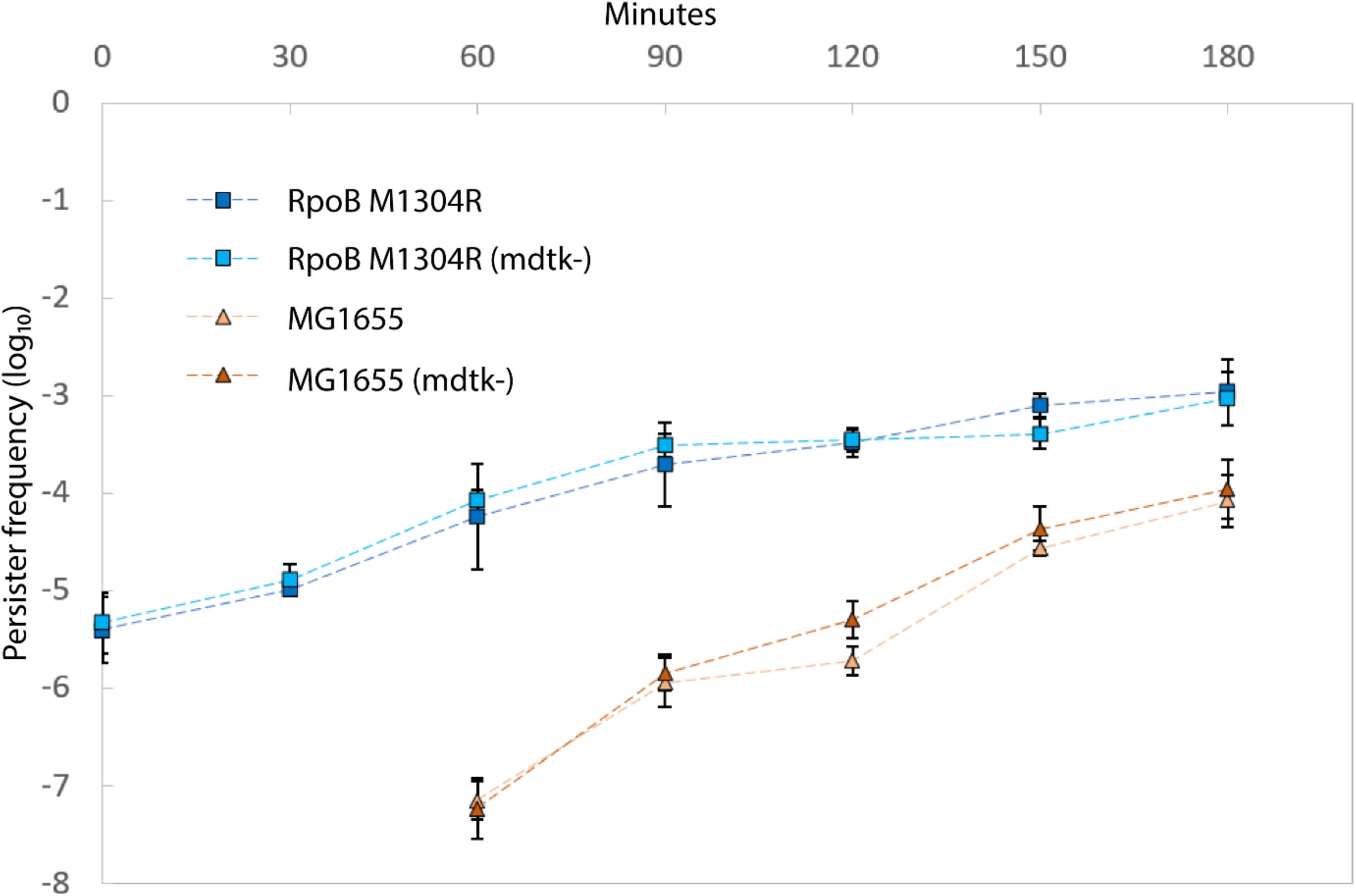
Deletion of *mdtK* has no effect on persistence. Comparison of persistence between MG1655 RpoB M1304R / MG1655 RpoB M1304R Δ*mdtk* (●) and MG1655 / MG1655 Δ*mdtk* (●) along the growth curve.

**Supplementary table 1:** Antibiotics

| Antibiotic | Cat # | Manufacturer | Concentration of stock | Stock age | Final dilution |
| --- | --- | --- | --- | --- | --- |
| Ampicillin | BP176025 | Fisher BioReagents | 100mg/ml (water) | ≤ 1 months (-20C) or 4 freeze cycles | 100mg/L |
| Kanamycin | 100218-998 | VWR Lifescience | 50mg/ml (water) | ≤ 3 months (-20C) | 50mg/L |
| Chloramphenicol | TCC2255 | TCI | 25 mg/ml (70% ethanol) | ≤ 2months (-20C) | 0.63mg/L<br>25mg/L |
| Rifampicin | R3501 | Sigma-Aldrich | 50mg/ml (methanol) | ≤ 2months (-20C) | 100mg/L |
| Ciprofloxacin | NC9553331 | Lkt Laboratories | 20, 10, 1, 0.1, 0.01mg/ml (water) | ≤ 1 months (-20C) or 4 freeze cycles | 1mg/L or 4mg/L |

**Supplementary table 2:** Strains

| Name | Relevant amino acid substitution | Genotype | Source |
| --- | --- | --- | --- |
| MG1655 | - | F<sup>-</sup>, lambda<sup>-</sup>, rph-1 | Plunkett, G. III |
| OKR5 | RpoB L533P | MG1655, *rpoB*1598 T>C | This study |
| OKR13 | RpoB M1304R | MG1655, *rpoB*3911 T>G, Δ*AceB* | This study |
| MG1655, pDJJ11 |  | MG1655, pDJJ11 | This study |
| OKR5, pDJJ11 | RpoB L533P | MG1655, *rpoB*1598 T>C, pDJJ11 | This study |
| OKR13, pDJJ11 | RpoB M1304R | MG1655, *rpoB*3911 T>G, Δ*AceB*, pDJJ11 | This study |
| OKR1 | GyrB S464Y | MG1655, *gyrB*1391 C>A | This study |
| OKR1, pUA66-M | GyrB S464Y | MG1655, *gyrB*1391 C>A, pUA66 (*mdtK*-*p::GFP*, *kanR*) | This study |
| OKR1, pUA66-I | GyrB S464Y | MG1655, *gyrB*1391 C>A, pUA66 (*ilvL*-*p::GFP*, *kanR*) | This study |
| OKR1, pUA66-H | GyrB S464Y | MG1655, *gyrB*1391 C>A, pUA66 (*hisL*-*p::GFP*, *kanR*) | This study |
| OKR2, pUA66-M | RpoB M1304R, GyrB S464Y | MG1655, *gyrB*1391 C>A, *rpoB*3911 T>G, pUA66 (*mdtK*:*GFP*, *kanR*) | This study |
| OKR2, pUA66-I | RpoB M1304R, GyrB S464Y | MG1655, *gyrB*1391 C>A, *rpoB*3911 T>G, pUA66 (*ilvL*:*GFP*, *kanR*) | This study |
| OKR2, pUA66-H | RpoB M1304R, GyrB S464Y | MG1655, *gyrB*1391 C>A, *rpoB*3911 T>G, pUA66 (*hisL*:*GFP*, *kanR*) | This study |
| OKR2 | RpoB M1304R, GyrB S464Y | MG1655, *gyrB*1391 C>A, *rpoB*3911 T>G | This study |
| OKR3 | RpoB E1279A, GyrB S464Y | MG1655, *gyrB*1391 C>A, *rpoB*3836 A>C | This study |
| OKR4 | RpoB V1275G, GyrB S464Y | MG1655, *gyrB*1391 C>A, *rpoB*3824 T>G:*kanR* | This study |
| MG1655(p) | - | Δ*dgoR*, Δ*relA*, Δ*spoT* | This study |
| OKR5(p) | RpoB L533P | MG1655, Δ*dgoR*, Δ*relA*, Δ*spoT* Δ*aceB*:*kanR*, *rpoB*1598 T>C | This study |
| OKR13(p) | RpoB M1304R | MG1655, Δ*dgoR*, Δ*relA*, Δ*spoT* Δ*aceB*:*kanR*, *rpoB*3911 T>G | This study |
| OKR1(p) | GyrB S464Y | MG1655, *gyrB*1391 C>A, Δ*relA*, Δ*spoT* | This study |
| OKR15(p) | GyrB S464Y, RpoB L533P | MG1655, *gyrB*1391 C>A, *ΔrelA*, *ΔspoT*, *ΔAceB:kanR*, *rpoB*1598 T>C | This study |
| OKR113(p) | GyrB S464Y, RpoB M1304R | MG1655, *gyrB*1391 C>A, *rpoB*3911 T>G, *ΔrelA*, *ΔspoT* | This study |
| OKR1135(p) | GyrB S464Y, RpoB M1304R, RpoB L533P | MG1655, *gyrB*1391 C>A, *rpoB*3911 T>G, *ΔrelA*, *ΔspoT*, *rpoB*1598 T>C | This study |
| OKR6 | GyrA S83L | MG1655, *gyrA*248 C>T | This study |
| OKR65 | RpoB L533P, GyrA S83L | MG1655, *rpoB*1598 T>C, *gyrA*248 C>T | This study |
| OKR613 | RpoB M1304R, GyrA S83L | MG1655, *rpoB*3911 T>G, *gyrA*248 C>T, *ΔAceB:kanR* | This study |
| WCUH29 | - | HA-MRSA | (82) |
| WCUH29-5 | RpoBS486L | *rpoB*1457 C>T | This study |

**Supplementary table 3:**
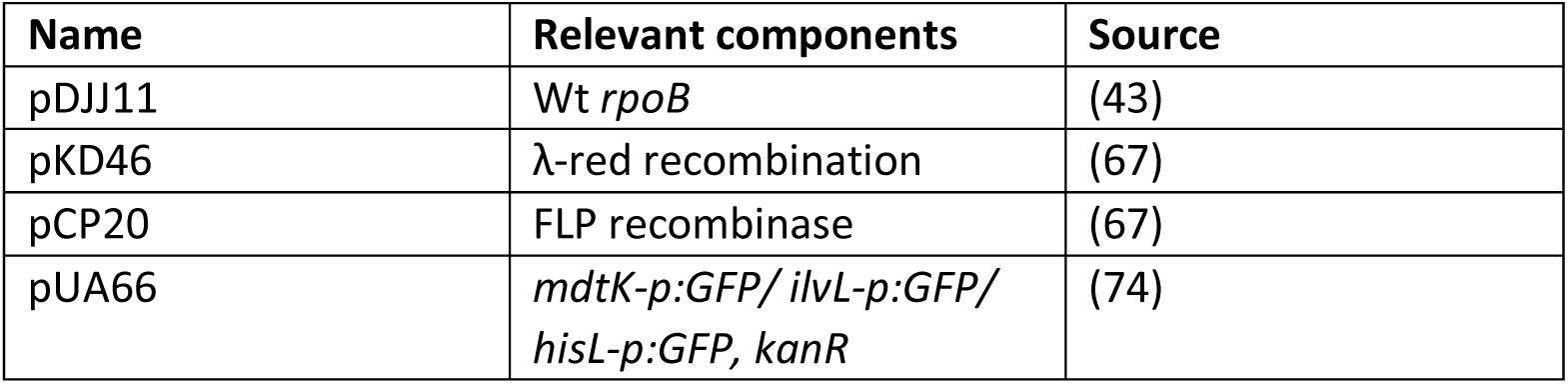
Plasmids

| **Name** | **Relevant components** | **Source** |
| --- | --- | --- |
| pDJJ11 | Wt *rpoB* | (43) |
| pKD46 | λ-red recombination | (67) |
| pCP20 | FLP recombinase | (67) |
| pUA66 | *mdtK-p:GFP/ ilvL-p:GFP/*<br>*hisL-p:GFP, kanR* | (74) |

**Supplementary table 4:**
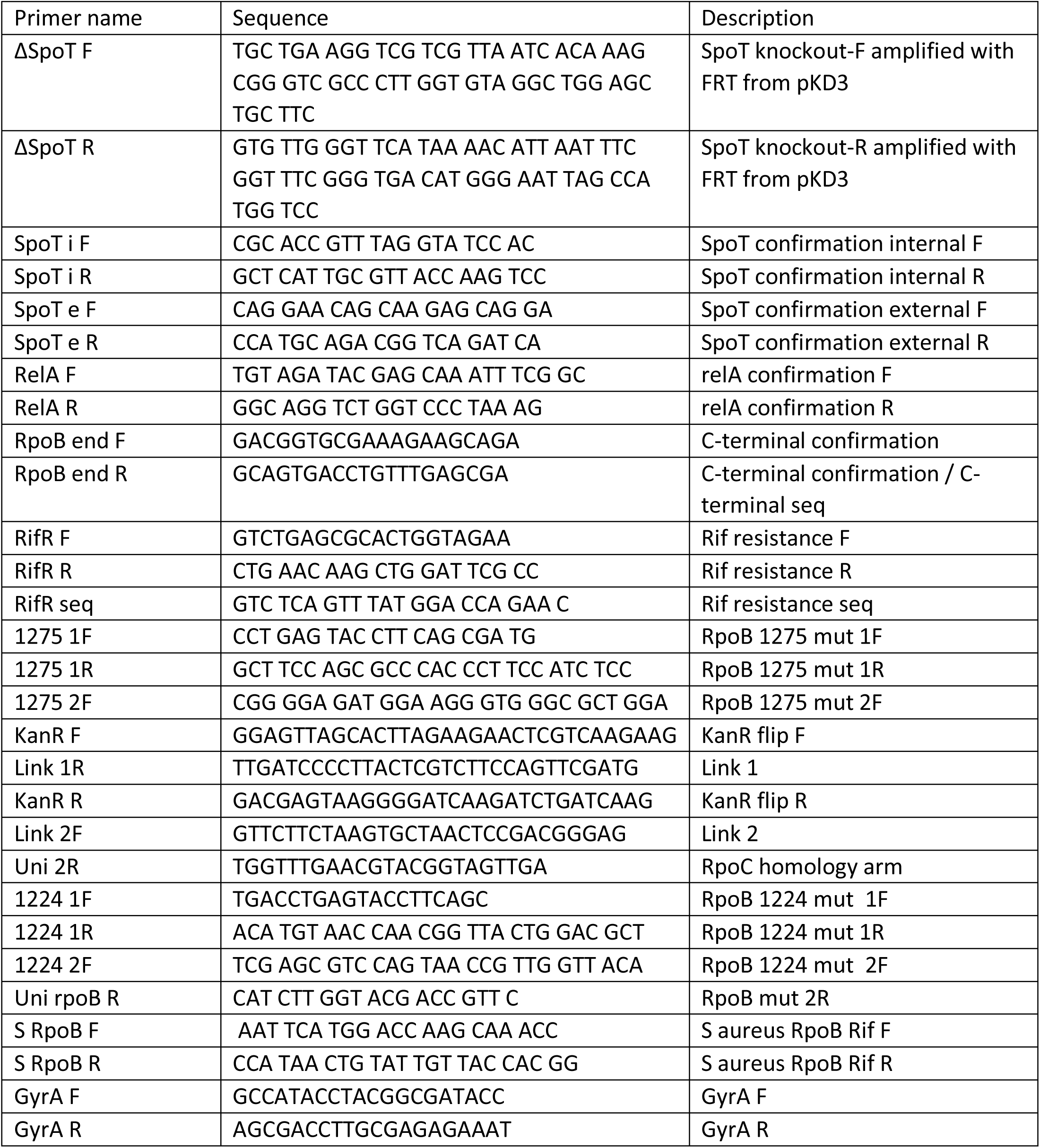
Primers

**Supplementary File 1:**
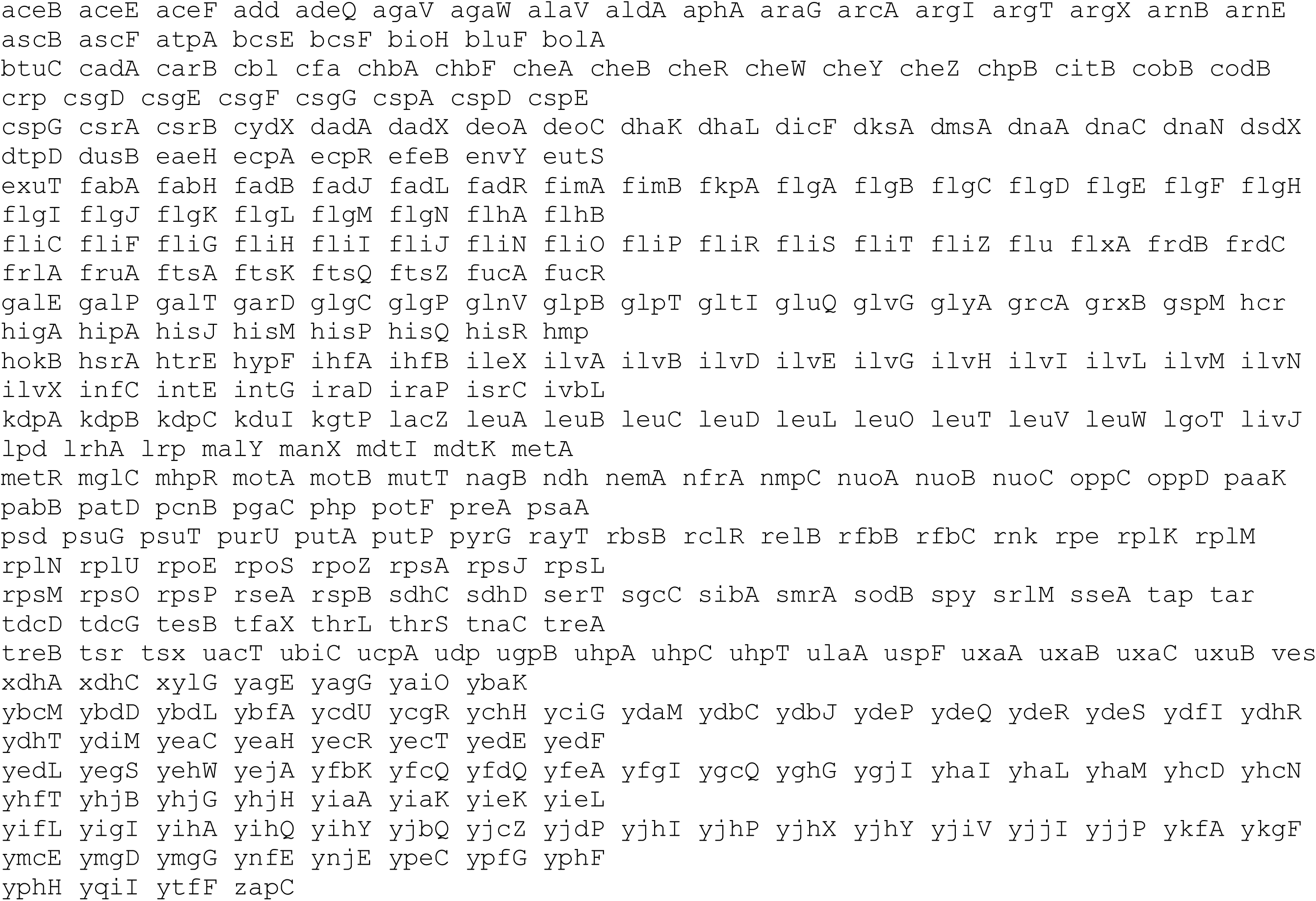
DE genes between RpoB M1304R and its parent; gene names are organized by row

**Supplementary File 2:**
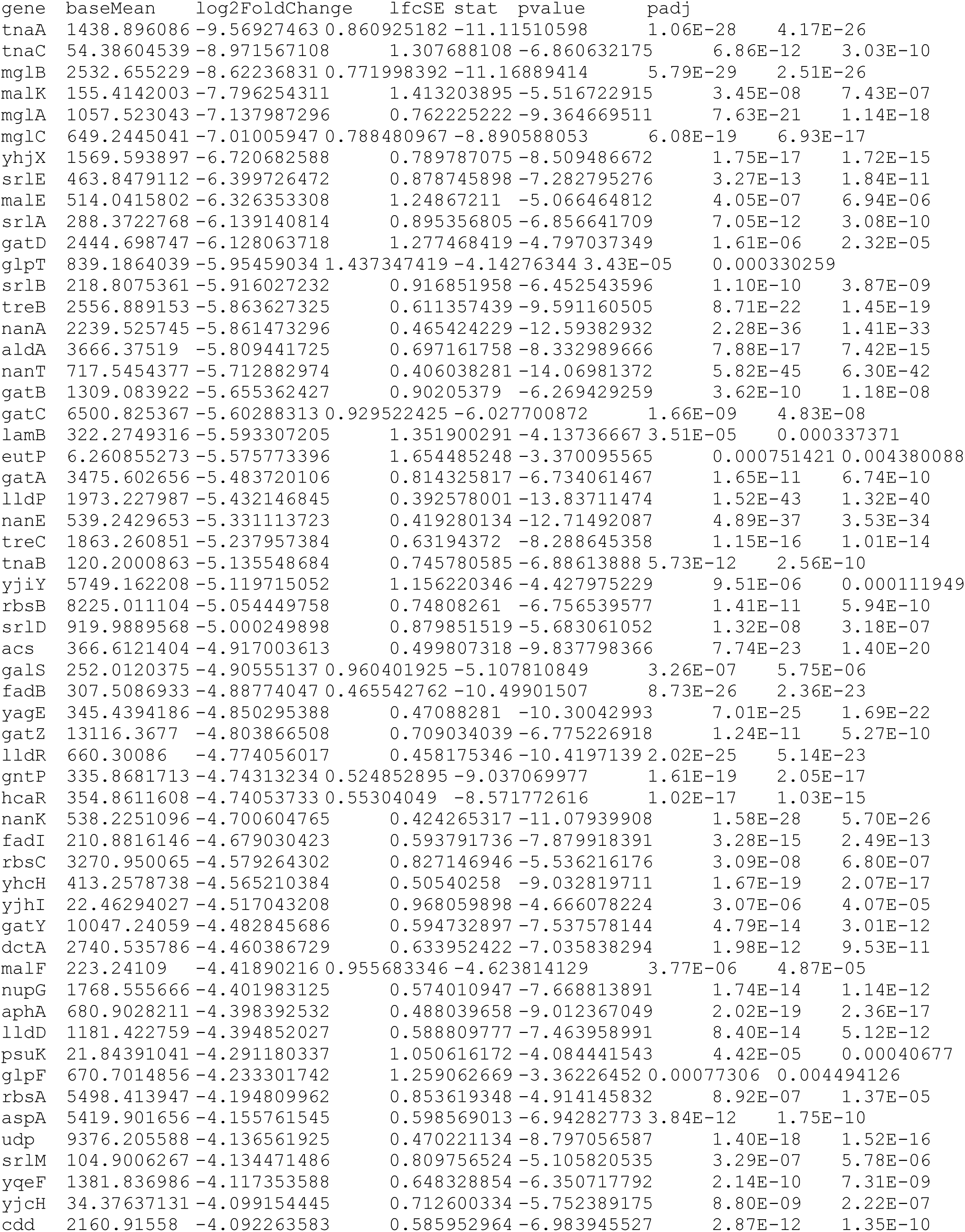
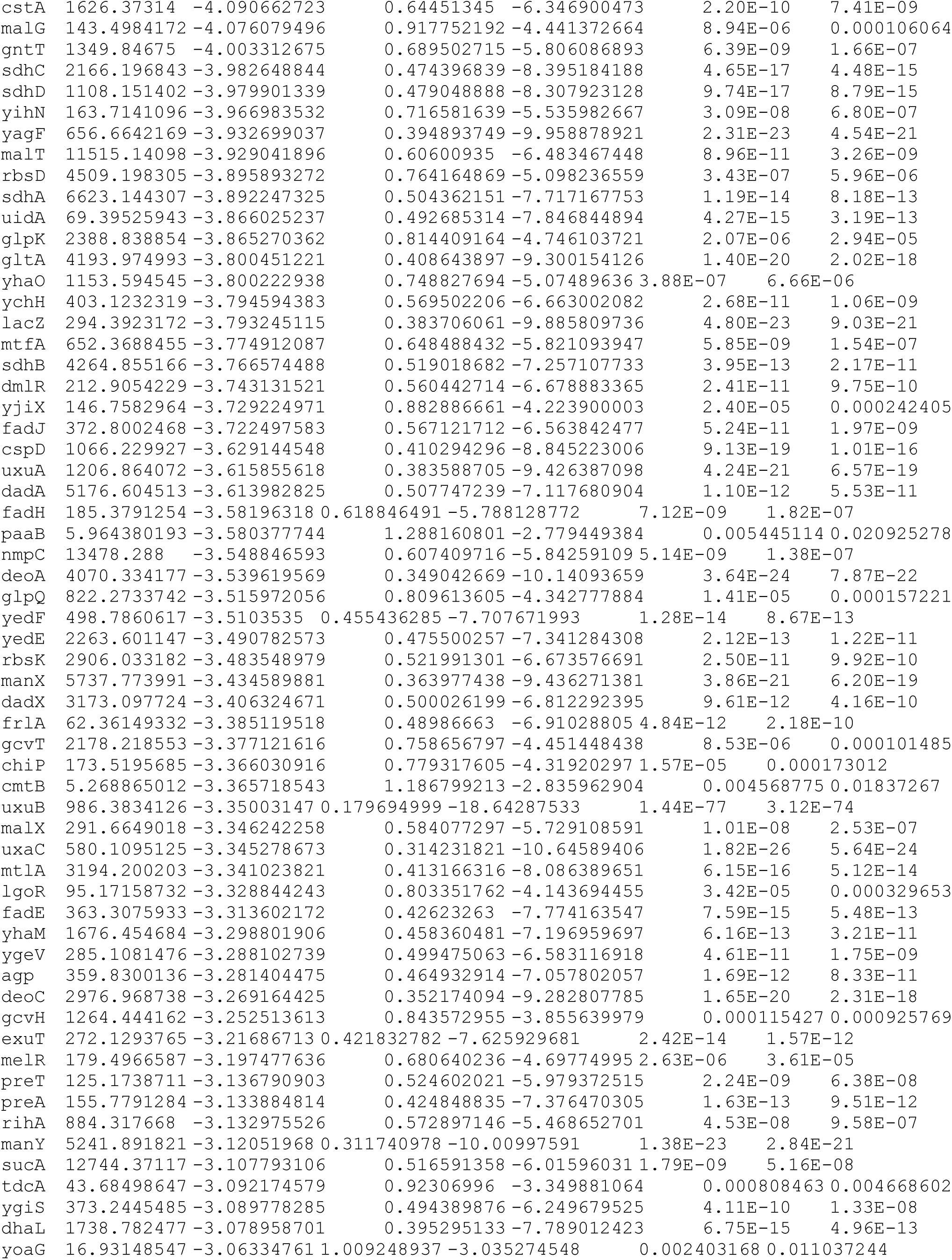
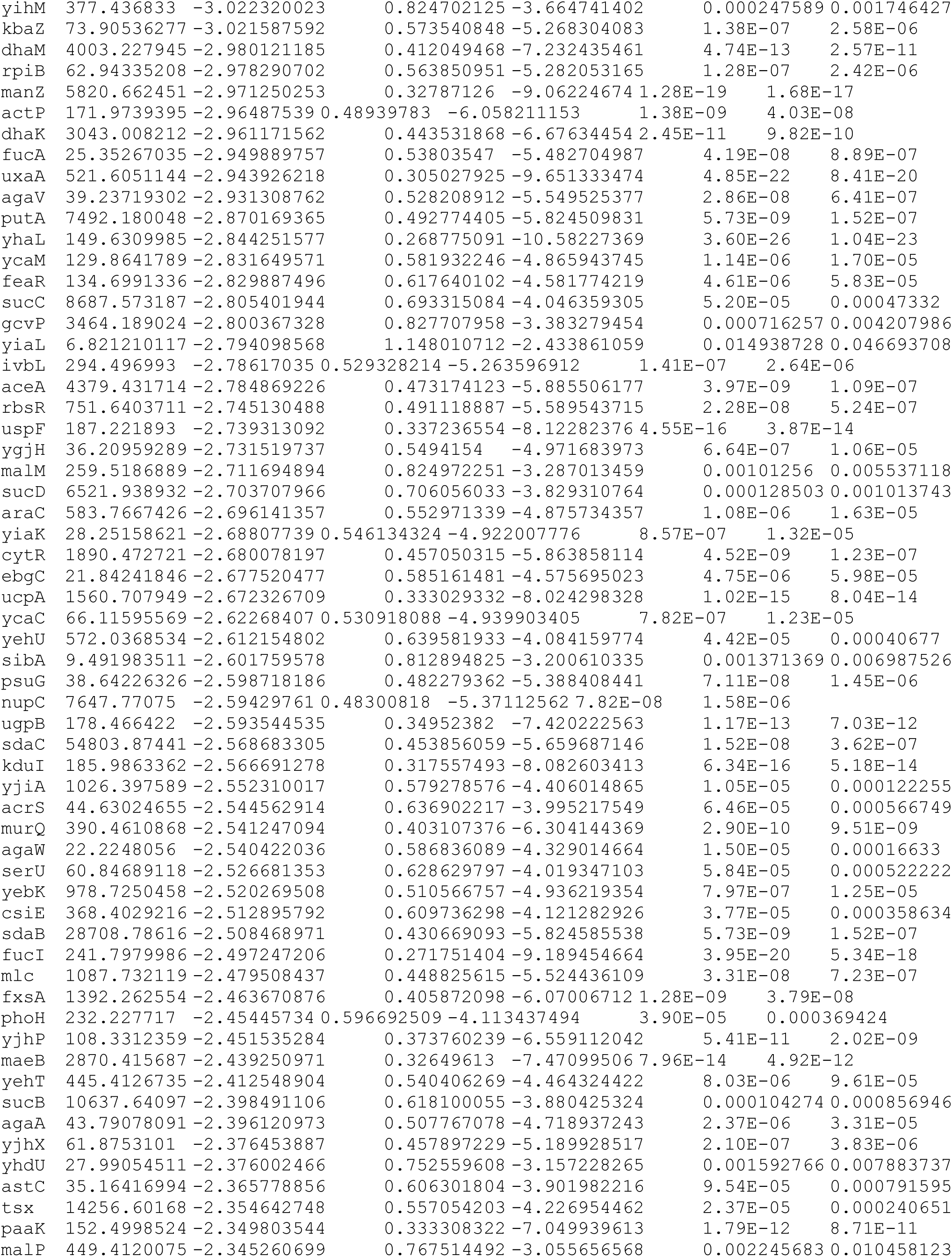
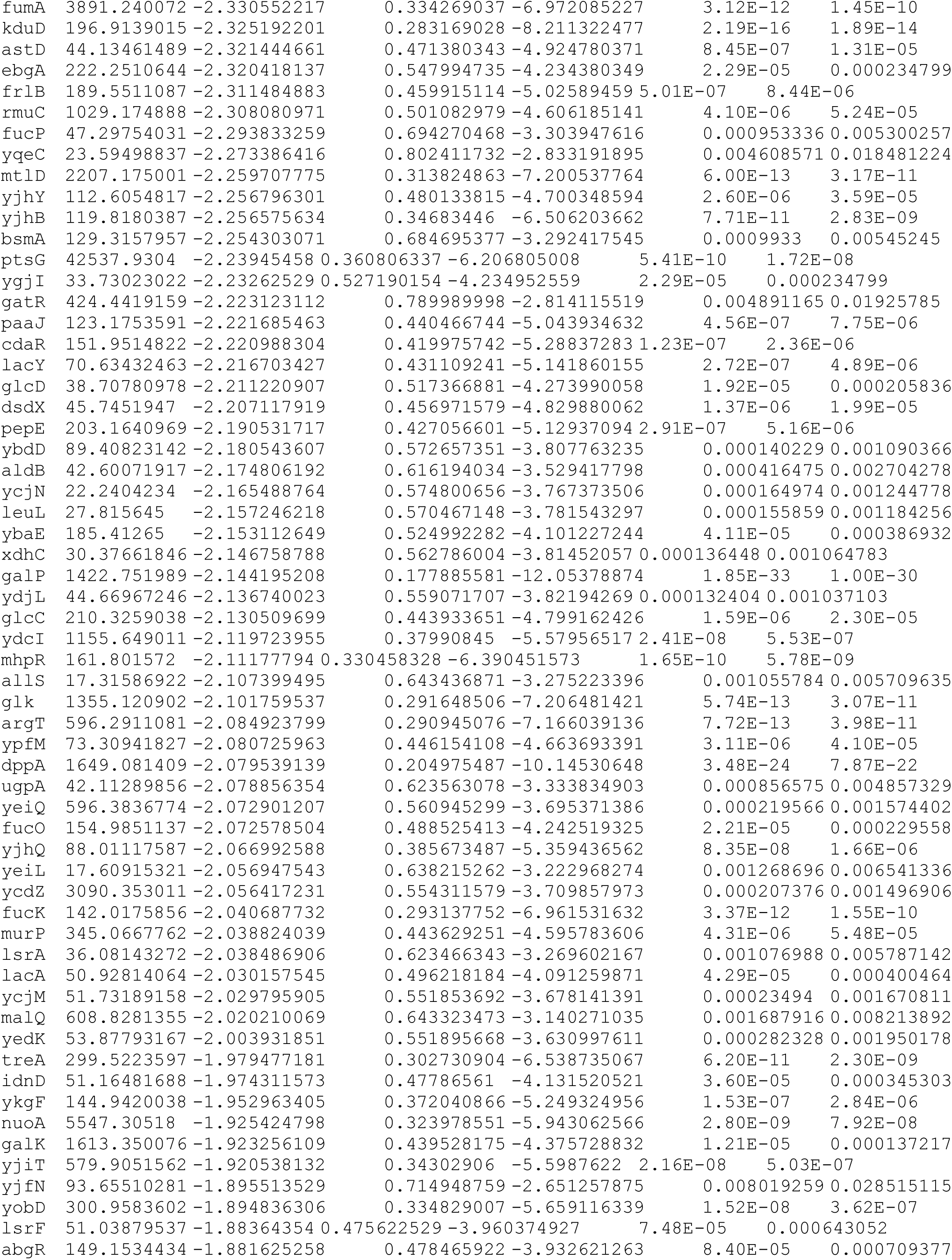
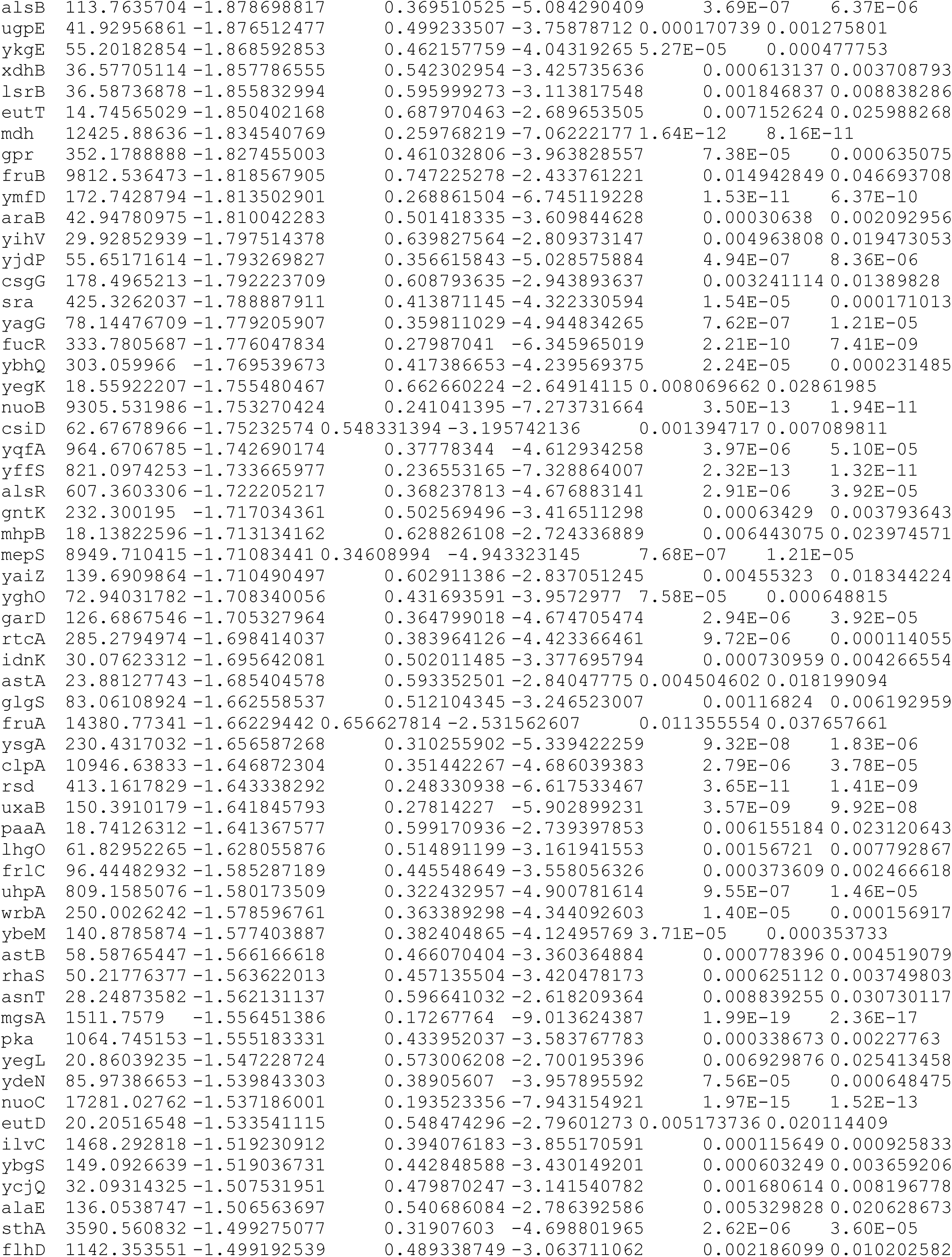
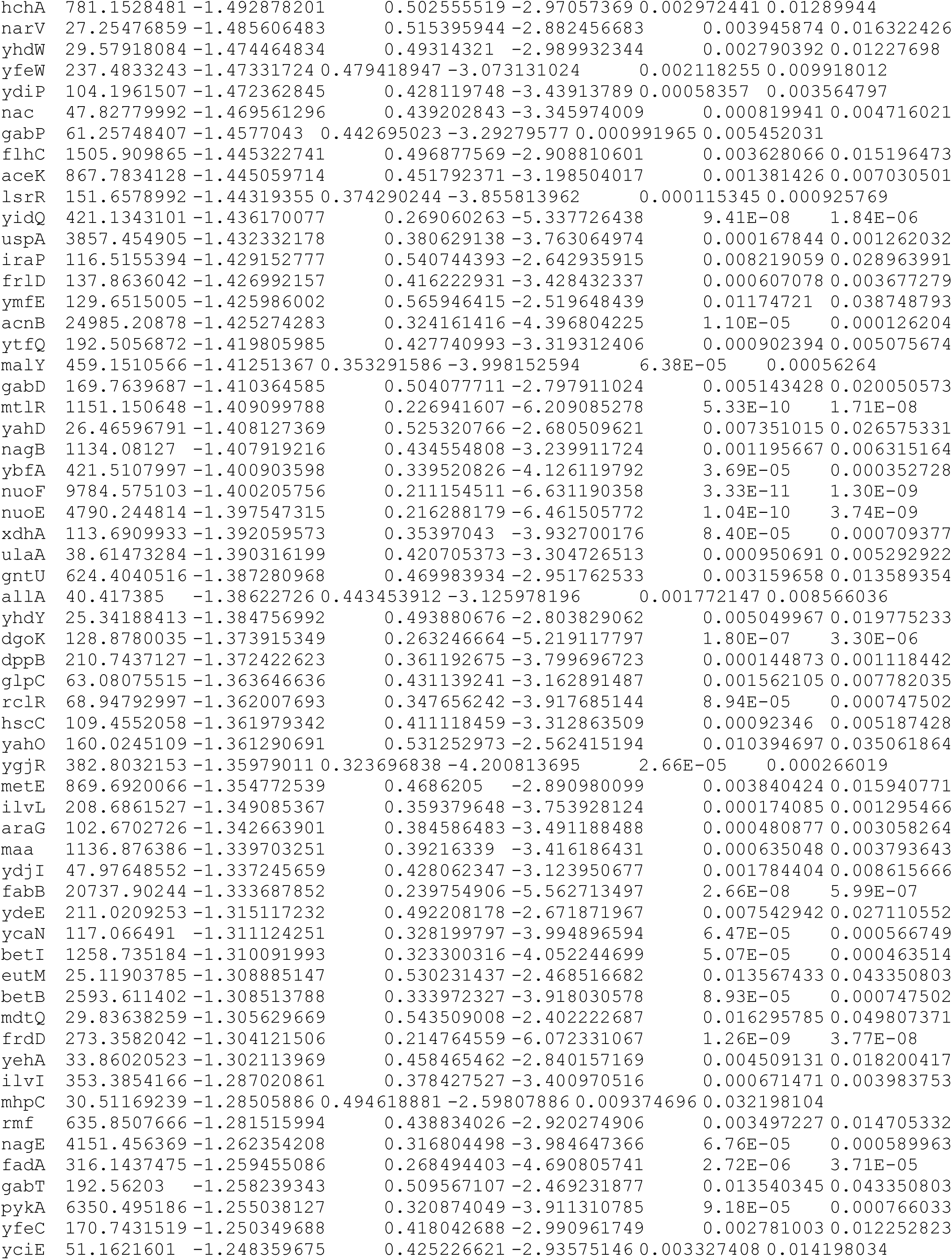
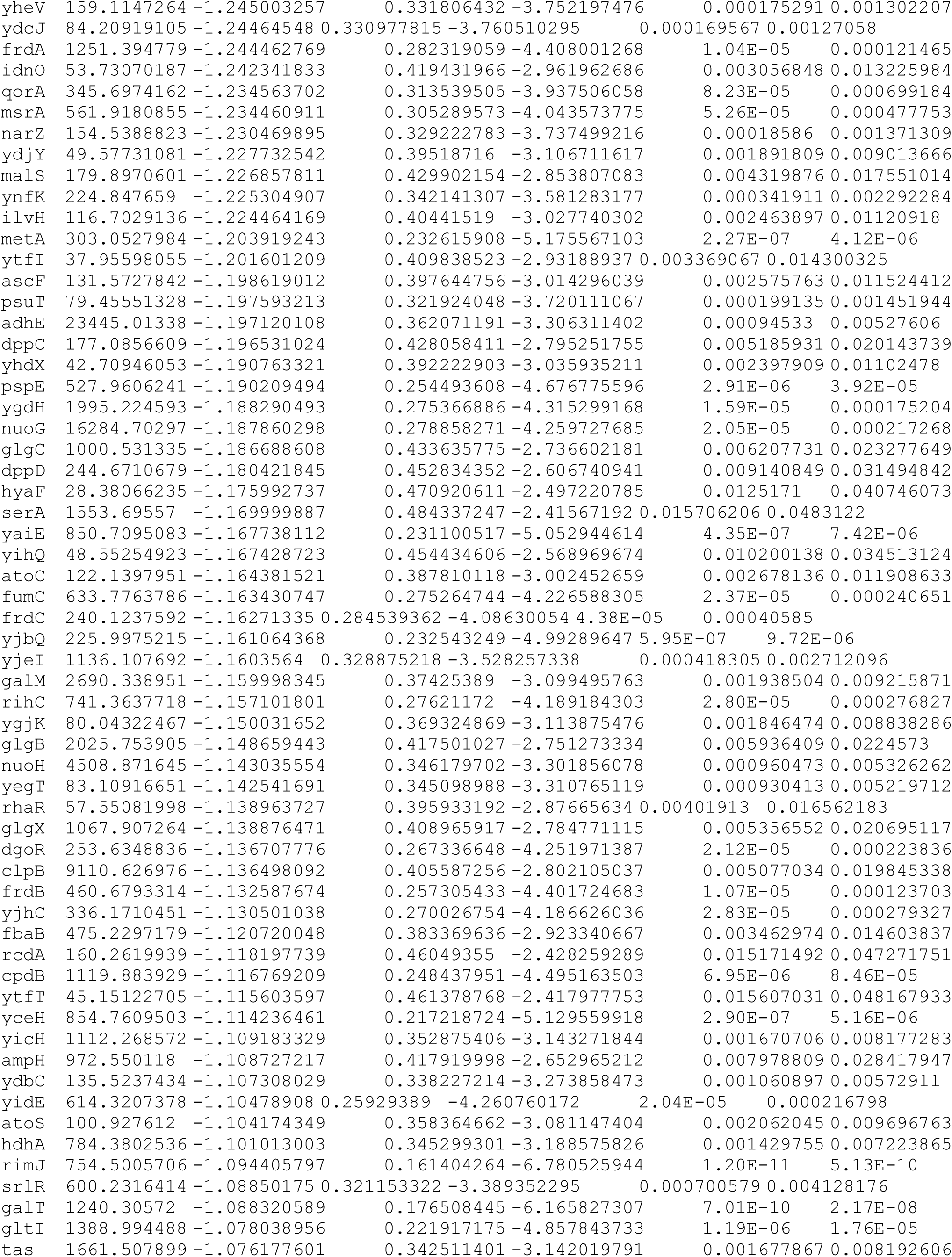
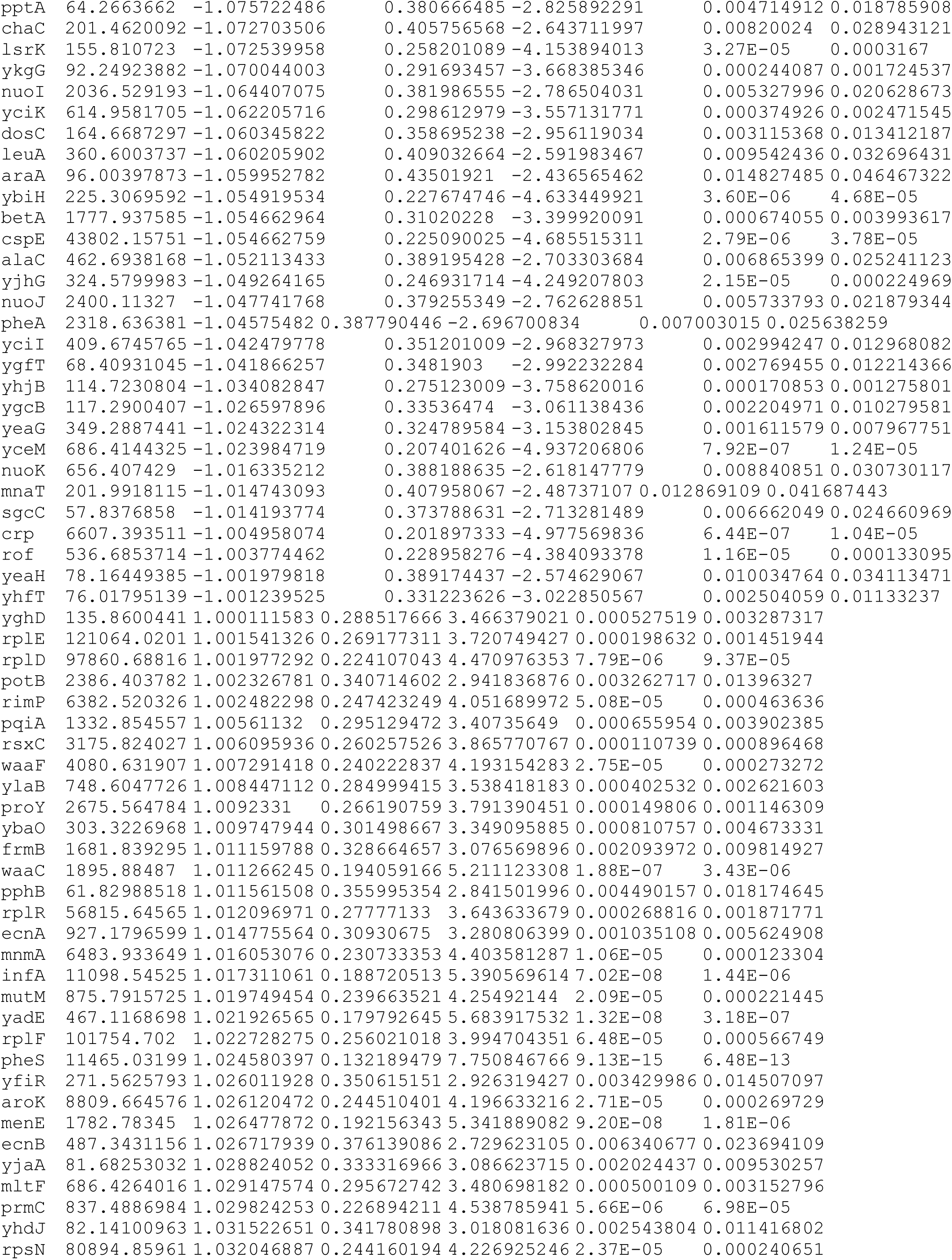
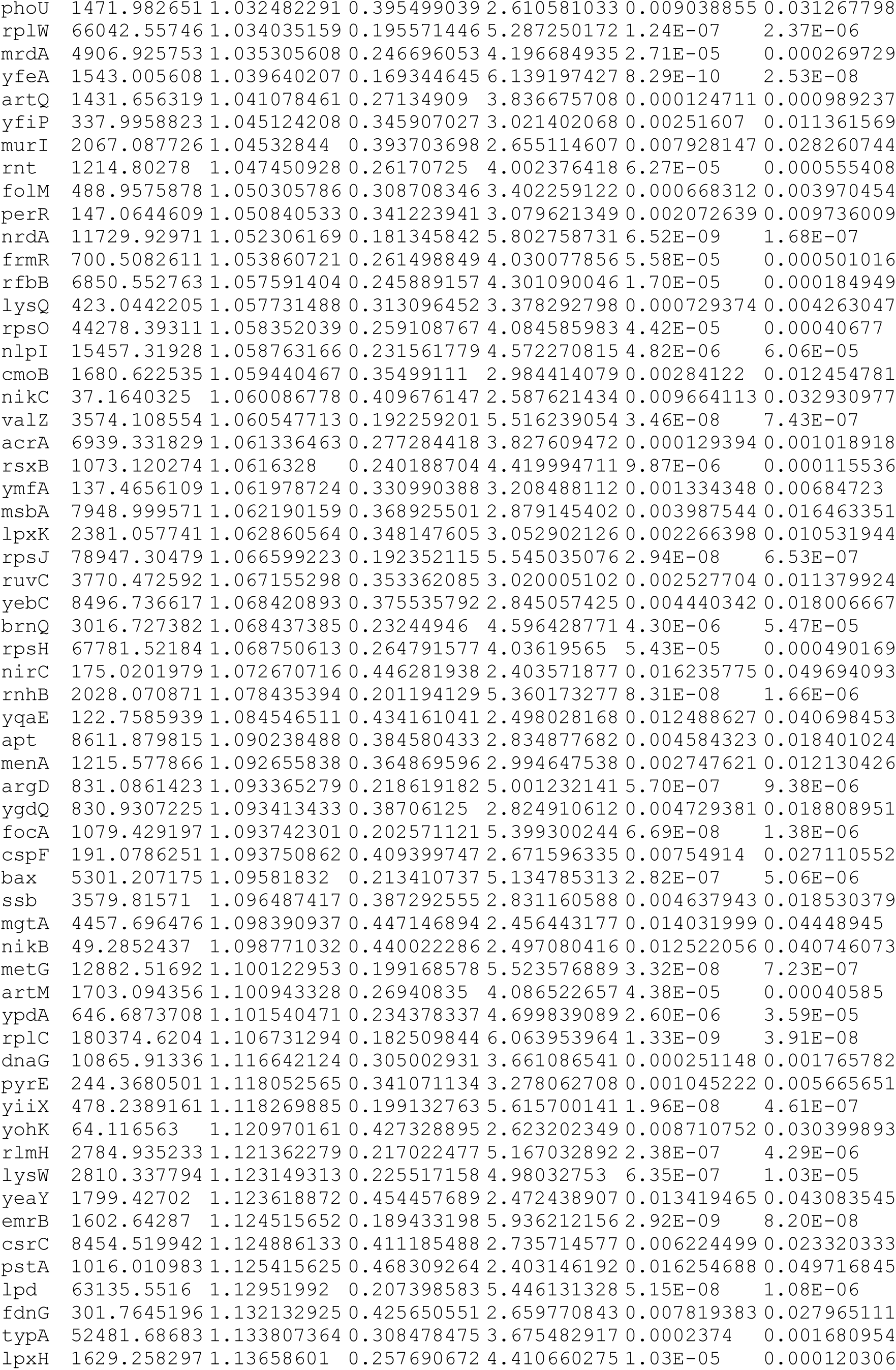
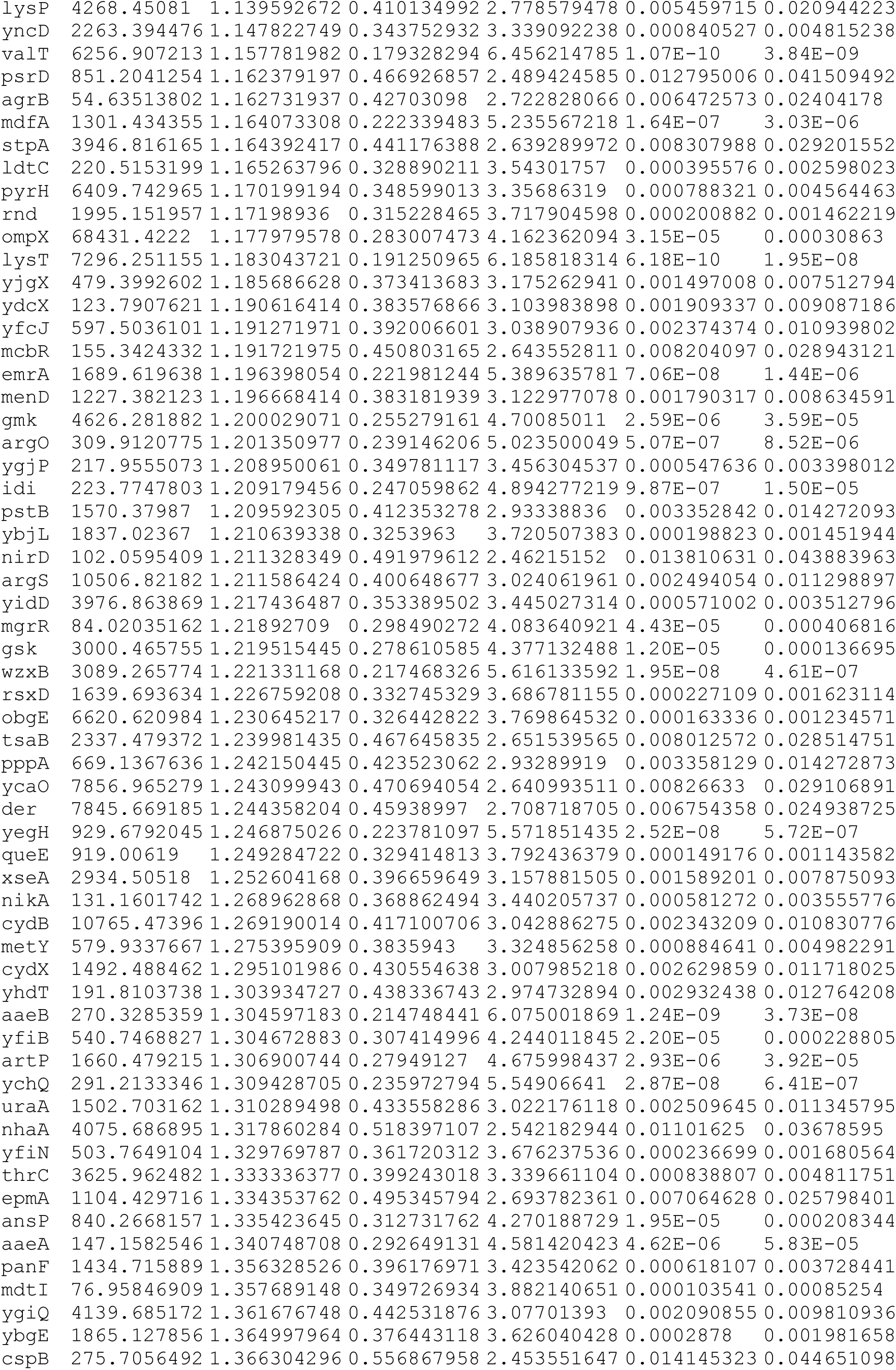
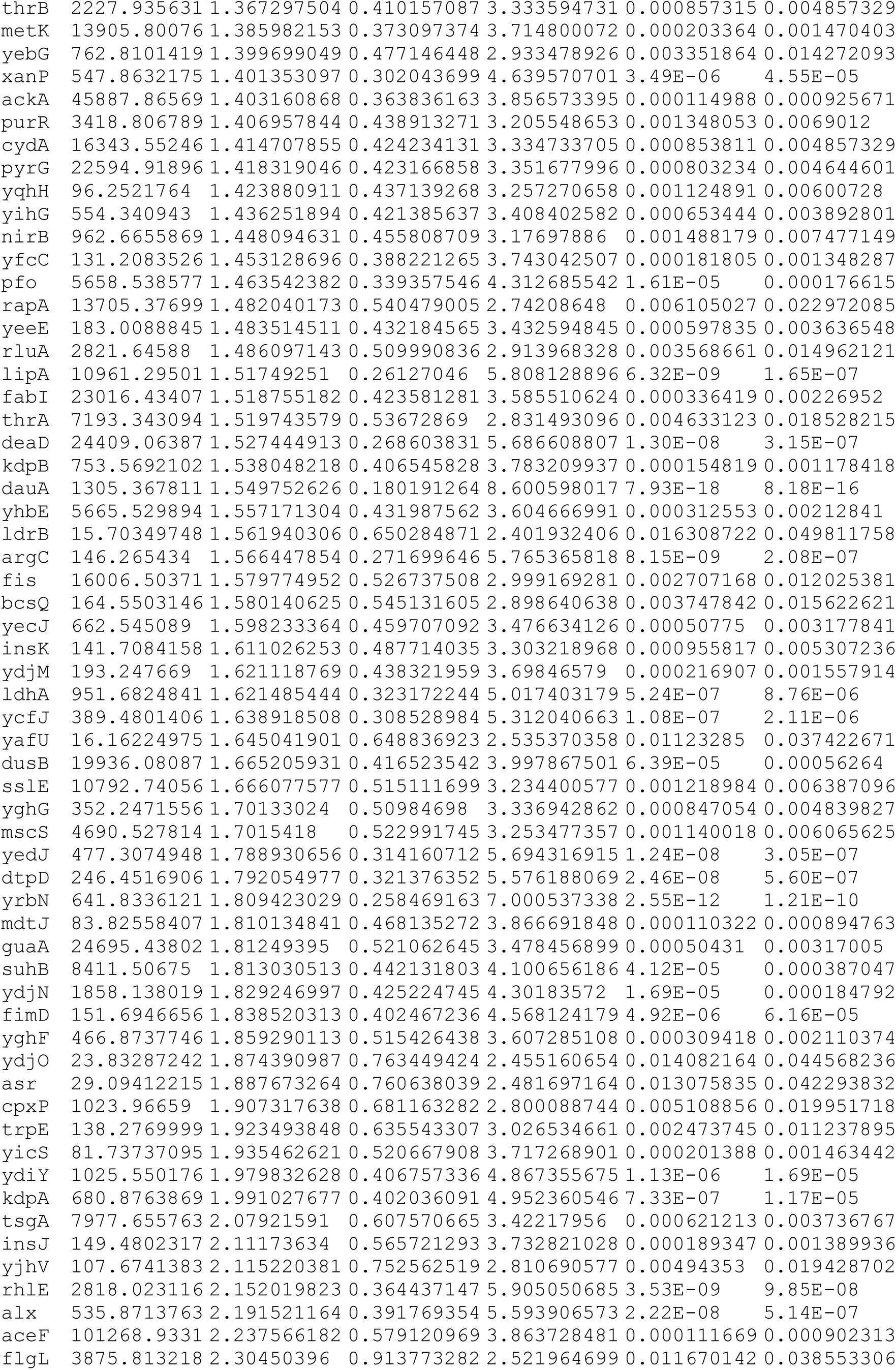
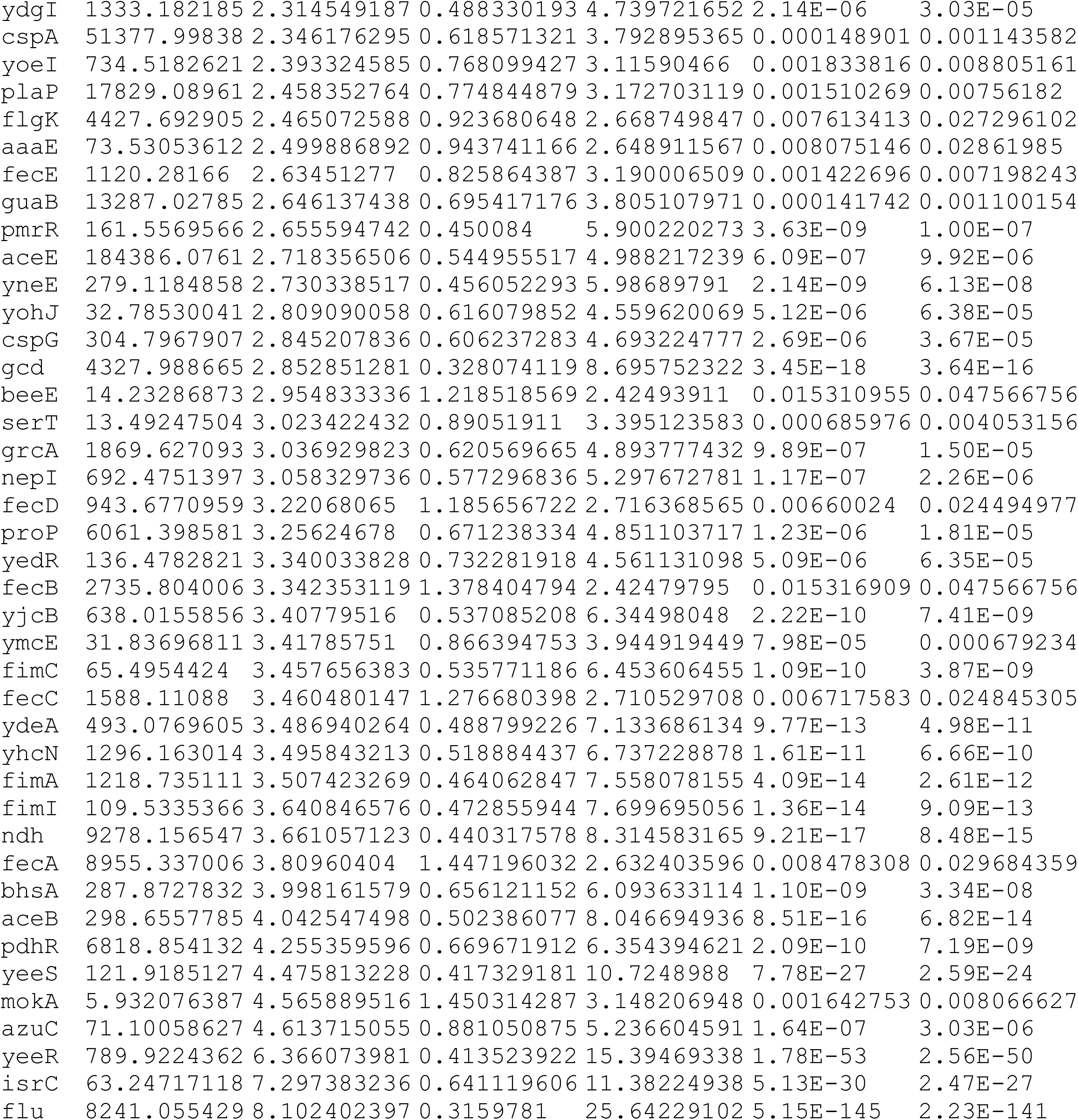
DE genes in both RpoB L533P and RpoB M1304R mutants relative to wt

**Supplementary File 3 (only top 50 genes are shown):**
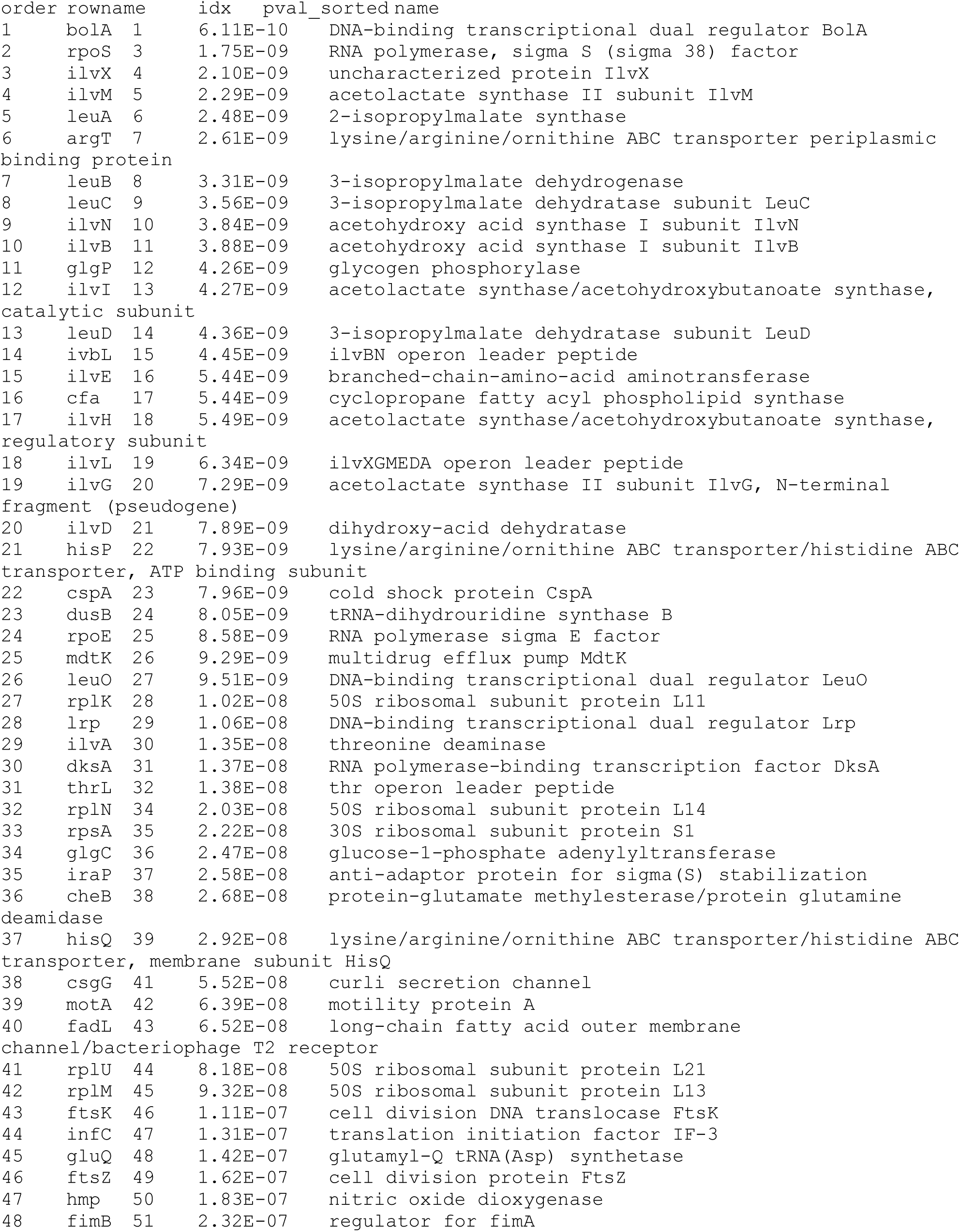
Genes, their functions, and probabilities of rejecting the null hypothesis of non-differential expression between RpoB M1304R and wt

## References

1. Nguyen D, Joshi-Datar A, Lepine F, Bauerle E, Olakanmi O, Beer K, Singh PK. 2011. Active starvation responses mediate antibiotic tolerance in biofilms and nutrient-limited bacteria. Science 334(6058):982–986.

2. Dörr T, Lewis K, Vulić M. 2009. SOS response induces persistence to fluoroquinolones in Escherichia coli. PLoS Genetics 5(12): e1000760.

3. Leung V, Lévesque CM. 2012. A stress-inducible quorum-sensing peptide mediates the formation of persister cells with noninherited multidrug tolerance. Journal of Bacteriology 194(9): 2265–2274.

4. Hu M, Zhang C, Mu Y, Shen Q, Feng Y. 2010. Indole affects biofilm formation in bacteria. Indian Journal of Microbiology 50(4):362–368.

5. Vega N, Allison K, Khalil A, Collins J. 2012. Signaling-mediated bacterial persister formation. Nature Chemical Biology 8(5): 431–433.

6. Wu N, He L, Cui P, Zhang Y. 2015. Ranking of persister genes in the same Escherichia coli genetic background demonstrates varying importance of individual persister genes in tolerance to different antibiotics. Frontiers of Microbiology 30(6):1003.

7. Chant E, Summers D. 2007. Indole signaling contributes to the stable maintenance of Escherichia coli multicopy plasmids. Molecular Microbiology 63(1):35–43.

8. Gefen O, Balaban NQ. 2009. The importance of being persistent: Heterogeneity of bacterial populations under antibiotic stress: Review article. FEMS Microbiology Reviews 33(4):704–717.

9. Rotem E, Lionger A, Ronin I, Balaban NQ. 2010. Regulation of phenotypic variability by a threshold-based mechanism underlies bacterial persistence. Proceedings of the National Academy of Sciences of the United States of America 107(28):12541–12546.

10. Moyed HS, Bertrand K. 1983. hipA, a newly recognized gene of Escherichia coli K-12 that affects frequency of persistence after inhibition of murein synthesis. Journal of Bacteriology 155(2): 768–775.

11. Moyed HS, Broderick SH. 1986. Molecular cloning and expression of hipA, a gene of Escherichia coli K-12 that affects frequency of persistence after inhibition of murein synthesis. Journal of Bacteriology 166(2):399–403.

12. Keren I, Shah D, Spoering A, Kaldalu N, Lewis K. 2004. Specialized persister cells and the mechanism of multidrug tolerance in Escherichia coli. Journal of Bacteriology 186(24):8172–8180.

13. Tripathi A, Dewan P, Siddique S, Varadarajan R. 2014. MazF-induced growth inhibition and persister generation in *Escherichia coli*. Journal of Biological Chemistry 289(7):4191–4205.

14. Shah D, Zhang Z, Khodursky A, Lewis K. 2006. Persisters: a distinct physiological state of *E. coli*. BMC Microbiology 6(1):53.

15. Braeken K, Moris M, Daniels R, Vanderleyden J, Michiels J. 2006. New horizons for (p)ppGpp in bacterial and plant physiology. Trends in Microbiology 14(1):45–54.

16. Amato S, Orman A, Brynildsen M. 2013. Metabolic control of persister formation in Escherichia coli. Molecular Cell 50(4):475–487.

17. Cashel M. 1970. Inhibition of RNA polymerase by ppGpp, a nucleotide accumulated during the stringent response to amino acid starvation in *E. coli*. Cold Spring Harbor Symposium Quantitative Biology 35: 407–413.

18. Wendrich TM, Blaha G, Wilson DN, Marahiel MA, Nierhaus KH. 2002. Dissection of the mechanism for the stringent factor RelA. Molecular Cell 10(4):779–788.

19. Xiao H, Kalman M, Ikehara K, Cashel M. 1991. Residual guanosine 3’,5’-bispyrophosphate synthetic activity of relA null mutants can be eliminated by spoT null mutations. The Journal of Biological Chemistry, 266(9), 5980–5990.

20. Traxler MF, Summers SM, Nguyen HT, Zacharia VM, Hightower GA, Smith JT, Conway T. 2008. The global, ppGpp-mediated stringent response to amino acid starvation in Escherichia coli. Molecular Microbiology, 68(5), 1128–1148.

21. Kuroda A, Tanaka S, Ikeda T, Kato J, Takiguchi N, Ohtake H. 1999. Inorganic polyphosphate kinase is required to stimulate protein degradation and for adaptation to amino acid starvation in Escherichia coli. Proceedings of the National Academy of Sciences of the United States of America, 96(25), 14264–14269.

22. Monode J. 1949. The growth of bacterial culture. Annual Reviews 3:371–394.

23. Korch SB, Henderson TA, Hill TM. 2003. Characterization of the hipA7 allele of Escherichia coli and evidence that high persistence is governed by (p)ppGpp synthesis. Mol Microbiol 50(4):1199–1213.

24. Yamaguchi Y, Inouye M. 2011. Regulation of growth and death in Escherichia coli by toxin– antitoxin systems. Nat Rev Microbiol 9:779–790.

25. Jin D, Gross C. 1989. Characterization of the pleiotropic phenotypes of rifampin-resistant rpoB mutants of Escherichia coli. Journal of Bacteriology 171(9):5229–5231.

26. Komp Lindgren P, Marcusson L, Sandvang D, Frimodt-Møller N, Hughes D. 2005. Biological cost of single and multiple norfloxacin resistance mutations in Escherichia coli implicated in urinary tract infections. Antimicrobial Agents and Chemotherapy 49(6):2343–2351.

27. Zhou Y, Jin D. 1998. The rpoB mutants destabilizing initiation complexes at stringently controlled promoters behave like “stringent” RNA polymerases in Escherichia coli. Proceedings of the National Academy of Sciences of the United States of America 95(6):2908–2913.

28. Mukhopadhyay J, Das K, Ismail S, et al. 2008. The RNA polymerase “Switch Region” is a target for inhibitors. Cell. 135(2):295–307.

29. Srivastava A, Talaue M, Liu S, et al. 2011. New target for inhibition of bacterial RNA polymerase: ‘switch region’. Current Opinions Microbiology. 14 (5):532–543.

30. Santos-Zavaleta A, Salgado H, Gama-Castro S, Callado-Vides J. 2019. RegulonDB v 10.5: tackling challenges to unify classic and high throughput knowledge of gene regulation in *E. coli* K-12. Nucleic Acids Research 47(6):212–220.

31. Keseler I, Mackie A, Santos-Zavaleta A, Karp P. 2017. The EcoCyc database: reflecting new knowledge about Escherichia coli K-12. Nucleic Acids Research 45(D1):543–550.

32. Koch A. 1971. The adaptive responses of Escherichia coli to a feast and famine existence. Advances in Microbial Physiology 6:147–217.

33. Kuroda A, Tanaka S, Ikeda T, Ohtake H. 1999. Inorganic polyphosphate kinase is required to stimulate protein degradation and for adaptation to amino acid starvation in Escherichia coli. Proceedings of the National Academy of Sciences of the United States of America 96(25):14264–14269.

34. Germain E, Guiraud P, Byrne D, Douzi B, Djendli M, Maisonneuve E. 2019. YtfK activates the stringent response by triggering the alarmone synthetase SpoT in Escherichia coli. Nature Communication. 10(1):1–12.

35. Little R, Ryals J. Bremer H. 1983. rpoB mutation in *Escherichia coli* alters control of ribosome synthesis by guanosine tetraphosphate. Journal of Bacteriology 154:787–792.

36. Potrykus K, Murphy H, Philippe N, Cashel M. 2011. ppGpp is the major source of growth rate control in *E. coli*. Environmental Microbiology 13(3):563–575.

37. Deris JB, Kim M, Zhang Z, Okano H, Hermsen R, Groisman A, Hwa T. 2013. The innate growth bistability and fitness landscapes of antibiotic-resistant bacteria. Science 342(6162):1237435.

38. Wattam A, Davis J, Assaf R, Stevens R. 2017. Improvements to PATRIC, the all-bacterial Bioinformatics Database and Analysis Resource Center. Nucleic Acids Research 45(D1):535–542.

39. Heddle J, Maxwell A. 2002. Quinolone-binding pocket of DNA gyrase: role of GyrB. Antimicrobial Agents Chemotherapy 46(6):1805–15.

40. Yoshida H, Bogaki M, Nakamura M, Nakamura S. 1990. Quinolone resistance-determining region in the DNA gyrase gyrA gene of Escherichia coli. Antimicrobial Agents and Chemotherapy 34(6):1271–1272.

41. Pietsch F, Bergman J, Brandis G, Hughes D. 2017. Ciprofloxacin selects for RNA polymerase mutations with pleiotropic antibiotic resistance effects. Journal of Antimicrobial Chemotherapy 72(1):75–84.

42. Ostrer L, Khodursky R, Johnson J, Hiasa H, Khodursky A. 2019. Analysis of mutational patterns in quinolone resistance-determining regions of GyrA and ParC of clinical isolates. International Journal of Antimicrobial Agents 53(3):318–324.

43. Jin D, Gross C. 1988. Mapping and sequencing of mutations in the Escherichia coli rpoB gene that lead to rifampicin resistance. Journal of Molecular Biology 202(1):45–58.

44. Yoshida H, Bogaki M, Nakamura M, Yamanaka L, Nakamura S. 1991. Quinolone resistance-determining region in the DNA gyrase gyrB gene of Escherichia coli. Antimicrobial Agents and Chemotherapy 35(8):1647–1650.

45. Fridman O, Goldberg A, Ronin I, Shoresh N, Balaban N. 2014. Optimization of lag time underlies antibiotic tolerance in evolved bacterial populations. Nature 513(7518):418–421.

46. Tuomanen E, Durack D, Tomasz A. 1986. Antibiotic tolerance among clinical isolates of bacteria. Antimicrobial Agents and Chemotherapy 30(4):521–527.

47. Van den Bergh B, Michiels J, Wenseleers T, Michiels J. 2016. Frequency of antibiotic application drives rapid evolutionary adaptation of Escherichia coli persistence. Nature Microbiology 1(5):16020.

48. Morita Y, Kodama K, Shiota S, Tsuchiya T. 1998. NorM, a putative multidrug efflux protein, of Vibrio parahaemolyticus and its homolog in Escherichia coli. Antimicrobial Agents and Chemotherapy 42(7):1778–1782.

49. Gong F, Ito K, Nakamura Y, Yanofsky C. 2001. The mechanism of tryptophan induction of tryptophanase operon expression: tryptophan inhibits release factor-mediated cleavage of TnaC-peptidyl-tRNA(Pro). Proceedings of the National Academy of Sciences of the United States of America 98(16):8997–9001.

50. Hopkins F, Cole S. 1903. A contribution to the chemistry of proteids: Part II. The constitution of tryptophan, and the action of bacteria upon it. The Journal of Physiology 29(4-5):451–466.

51. Lacour S, Landini P. 2004. SigmaS-dependent gene expression at the onset of stationary phase in Escherichia coli: function of sigmaS-dependent genes and identification of their promoter sequences. Journal of Bacteriology 186(21):7186–7195.

52. Prüss B, Nelms J, Park C, Wolfe A. 1994. Mutations in NADH: ubiquinone oxidoreductase of Escherichia coli affect growth on mixed amino acids. Journal of Bacteriology 176(8):2143–2150.

53. Halsey C, Lei S, Wax J, Fey P. 2017. Amino acid catabolism in *Staphylococcus aureus* and the function of carbon catabolite repression. MBio, 8(1):e01434–16.

54. Nuxoll A, Halouska S, Sadykov M, Fey P. 2012. CcpA regulates arginine biosynthesis in *Staphylococcus aureus* through repression of proline catabolism. PLoS Pathogens 8(11):e1003033.

55. Stanley N, Lazazzera B. 2004. Environmental signals and regulatory pathways that influence biofilm formation. Molecular Microbiology 52(4):L917–924.

56. Reed JM, Olson S, Brees DF, Griffin CE, Grove RA, Davis PJ, Somerville GA. 2018. Coordinated regulation of transcription by CcpA and the *Staphylococcus aureus* two-component system HptRS. PloS One 13(12):e0207161.

57. Seidl K, Goerke C, Mack D, Bishoff M. 2008. *Staphylococcus aureus* CcpA affects biofilm formation. Infection and Immunity 76(5):2044–2050.

58. Brandis G, Pietsch F, Alemayehu R, Hughes D. 2015. Comprehensive phenotypic characterization of rifampicin resistance mutations in *Salmonella* provides insight into the evolution of resistance in *Mycobacterium tuberculosis*. Journal of Antimicrobial Chemotherapy 70(3):680–685.

59. Campbell EA, Korzheva N, Mustaev A, Murakami K, Nair S, Goldfarb A, Darst SA. 2001. Structural mechanism for rifampicin inhibition of bacterial RNA polymerase. Cell 104(6):901–912.

60. Telenti A, Imboden P, Marchesi F, Cole S. 1993. Detection of rifampicin-resistance mutations in *Mycobacterium tuberculosis*. Lancet 341(8846):647–650.

61. Wichelhaus T, Schäfer V, Brade V, Böddinghaus B. 1999. Molecular characterization of rpoB mutations conferring cross-resistance to rifamycins on methicillin-resistant *Staphylococcus aureus*. Antimicrobial Agents and Chemotherapy 43(11):2813–2816.

62. Brandis G, Wrande M, Liljas L, Hughes D. 2012. Fitness-compensatory mutations in rifampicin-resistant RNA polymerase. Molecular Microbiology 85(1):142–151.

63. Forrest G, Tamura K. 2010. Rifampin combination therapy for nonmycobacterial infections. Clinical Microbiology Reviews 23(1):14–34.

64. Fisher R, Gollan B, Helaine S. 2017. Persistent bacterial infections and persister cells. Nature Reviews Microbiology 15(8):453–464.

65. Riley M, Abe T, Arnaud MB, Berlyn MKB. 2006. Escherichia coli K-12: a cooperatively developed annotation snapshot--2005. Nucleic Acids Research 34(1):1–9.

66. Baba T, Ara T, Hasegawa M, Mori H. 2006. Construction of Escherichia coli K-12 in-frame, single-gene knockout mutants: the Keio collection. Molecular Systems Biology 2(1):2006–2008.

67. Datsenko K, Wanner B. 2000. One-step inactivation of chromosomal genes in Escherichia coli K-12 using PCR products. Proceedings of the National Academy of Sciences of the United States of America 97(12):6640–6645.

68. Missiakas D, Schneewind O. 2013. Growth and laboratory maintenance of *Staphylococcus aureus*. Current Protocols in Microbiology Chapter 9, Unit 9C.1.

69. Wiegand I, Hilpert K, Hancock R. 2008. Agar and broth dilution methods to determine the minimal inhibitory concentration (MIC) of antimicrobial substances. Nature Protocols 3(2):163–175.

70. Li H, Handsaker B, Wysoker A, Durbin R. 2009. The Sequence Alignment/Map format and SAMtools. Bioinformatics 25(16):2078–2079.

71. Gutierrez A, Jain S, Bhargava P, Hamblin M, Lobritz MA, Collins JJ. 2017. Understanding and sensitizing density-dependent persistence to quinolone antibiotics. Molecular Cell 68(6):1147–1154.

72. Jensen K, Houlberg U, Nygaard P. 1979. Thin-layer chromatographic methods to isolate 32P-labeled 5-phosphoribosyl-alpha-1-pyrophosphate (PRPP): determination of cellular PRPP pools and assay of PRPP synthetase activity. Analytical Biochemistry 98(2):254–263.

73. Schindelin J, Arganda-Carreras I, Frise E, Cardona A. 2012. Fiji: an open-source platform for biological-image analysis. Nature Methods 9(7):676–682.

74. Zaslaver A, Bren A, Ronen M, Alon U. 2006. A comprehensive library of fluorescent transcriptional reporters for Escherichia coli. Nature Methods 3(8):623–628.

75. Pertea M, Kim D, Pertea G, Leek J, Salzberg S. 2016. Transcript-level expression analysis of RNA-seq experiments with HISAT, StringTie and Ballgown. Nature Protocols 11(9):1650–1667.

76. Liao Y, Smyth G, Shi W. 2019. The R package Rsubread is easier, faster, cheaper and better for alignment and quantification of RNA sequencing reads. Nucleic Acids Research, 47(8):e47.

77. Love M, Huber W, Anders S. 2014. Moderated estimation of fold change and dispersion for RNA-seq data with DESeq2. Genome Biology 15(12):550.

78. Langmead B. 2010. Aligning short sequencing reads with Bowtie. Current Protocols in Bioinformatics, chapter 11.

79. Langmead B, Salzberg S. 2012. Fast gapped-read alignment with Bowtie 2. Nature Methods 9(4):357–359.

80. Deatherage D, Barrick J. 2014. Identification of mutations in laboratory-evolved microbes from next-generation sequencing data using breseq. Methods in Molecular Biology 1151:165–188.

81. Quinlan A, Hall I. 2010. BEDTools: a flexible suite of utilities for comparing genomic features. Bioinformatics 26(6):841–842.

82. Lei T, Zhang Y, Yang J, Silverstein K, Ji Y. 2019. Complete Genome Sequence of Hospital-Acquired Methicillin-Resistant *Staphylococcus aureus* Strain WCUH29. Microbiology Resource Announcements 8(23):e00551–19.

